# Cerebral organoid model reveals excessive proliferation of human caudal late interneuron progenitors in Tuberous Sclerosis Complex

**DOI:** 10.1101/2020.02.27.967802

**Authors:** Oliver L. Eichmüller, Nina S. Corsini, Ábel Vértesy, Theresa Scholl, Victoria-Elisabeth Gruber, Angela M. Peer, Julia Chu, Maria Novatchkova, Mercedes F. Paredes, Martha Feucht, Jürgen A. Knoblich

## Abstract

Although the intricate and prolonged development of the human brain critically distinguishes it from other mammals^1^, our current understanding of neurodevelopmental diseases is largely based on work using animal models. Recent studies revealed that neural progenitors in the human brain are profoundly different from those found in rodent animal models^2–5^. Moreover, post-mortem studies revealed extensive migration of interneurons into the late-gestational and post-natal human prefrontal cortex that does not occur in rodents^6^. Here, we use cerebral organoids to show that overproduction of mid-gestational human interneurons causes Tuberous Sclerosis Complex (TSC), a severe neuro-developmental disorder associated with mutations in *TSC1* and *TSC2*. We identify a previously uncharacterized population of caudal late interneuron progenitors, the CLIP-cells. In organoids derived from patients carrying heterozygous *TSC2* mutations, dysregulation of mTOR signaling leads to CLIP-cell over-proliferation and formation of cortical tubers and subependymal tumors. Surprisingly, second-hit events resulting from copy-neutral loss-of-heterozygosity (cnLOH) are not causative for but occur during the progression of tumor lesions. Instead, EGFR signaling is required for tumor proliferation, opening up a promising approach to treat TSC lesions. Our study demonstrates that the analysis of developmental disorders in organoid models can lead to fundamental insights into human brain development and neuropsychiatric disorders.

---

Tuberous sclerosis complex (TSC) is a rare autosomal dominant disorder characterized by pathological malformations in multiple organs^7^. Among those, brain defects leading to severe neuropsychiatric symptoms like autism spectrum disorder (ASD), intractable seizures and intellectual disability (ID) are most debilitating and seen in the majority of patients^8^. Most patients have cortical tubers^9^, focal dysplastic regions in the cortex that are diagnosed by MRI and consist of dysmorphic neurons and giant cells. In addition, 80% of the patients display subependymal nodules (SEN) that form along the lateral ventricle and can develop into subependymal giant cell astrocytomas (SEGAs) in 10-15% of the patients^7,10^. It was thought that TSC pathogenesis is initiated by constitutive mTOR activity resulting from inactivation of the second allele^11^ along the lines of the classic Knudson two-hit hypothesis of tumorigenesis^12^. This is supported by existing mouse models and a spheroid model for TSC, as characteristic brain alterations are observed exclusively in *Tsc1* or *Tsc2* homozygous mutant mice and spheroids^13–18^. Genetic analysis in patients, however, revealed that loss of the second allele is frequent in SEN/SEGA, but rare in cortical tubers^19–22^, conflicting with the two-hit hypothesis. In addition, the cellular origins of cortical tubers and SEN/SEGAs remain unclear. Interestingly, recent work using human primary tissues suggested a common cell-of-origin for both lesions based on shared transcriptomic alterations^21^. In order to study the cell types and mechanisms leading to the generation of tumors and tubers we generated human cerebral organoids^23^ from patient-derived induced pluripotent stem cells (iPSCs) and compared our results to human primary material.

## Human cerebral organoids recapitulate TSC histopathology

To model the brain pathology of TSC, we derived human iPSCs from two patients with known *TSC2* mutations (Ext. Data Fig. 1). All selected patients have drug-resistant epilepsy and show cortical tubers and subependymal tumors (Ext. Data Fig. 1A-C). The first patient is germline mosaic, which allowed us to derive both *TSC2*^+/−^ and isogenic *TSC2*^+/+^ lines (Ext. Data Fig. 1D). For the second patient, isogenic controls were generated by scarless Crispr-based genome editing (Ext. Data Fig. 1E). To study proliferative phenotypes, organoids were cultured in a high-nutrient (H-) medium that promotes proliferation analogous to germinal areas that give rise to subependymal tumors in the brain (Fig. 1B). In order to mimic cortical tuber formation, we transitioned organoids to a low-nutrient (L-) medium that was designed by adapting a published formulation^24^ to 3D culture (Fig. 1B, see Materials and Methods for details).

**Figure 1.**
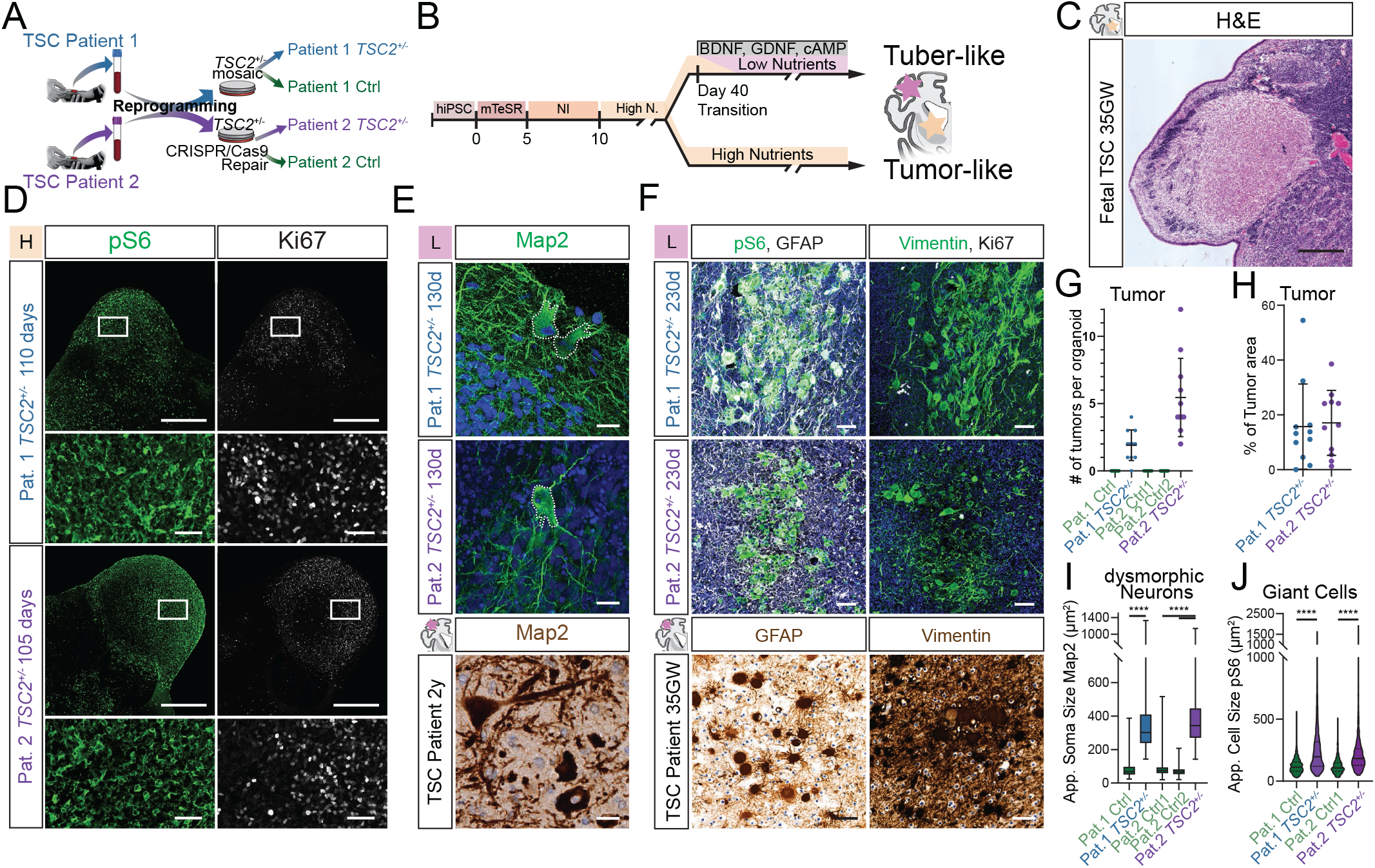
*TSC2*^+/−^-derived organoids recapitulate histopathology of TSC. **A.** Ctrl and *TSC2*^+/−^ cell lines were derived from two patients (see Materials and Methods for details). **B.** High- and Low-nutrition organoid protocols used to model distinct TSC phenotypes. **C.** H&E staining of 35GW fetal brain depicts histopathology of a fetal SEN. **D.** pS6 and Ki67 staining on 110- and 105 days-old *TSC2*^+/−^-derived organoids in high-nutrient medium identifies SEN-like structures. Lower panel shows higher magnification of inset. **E.** Map2 staining on 130 days-old organoids in L-medium shows dysmorphic neurons (top), with comparable morphology to those in a resected tuber of a 2 years-old patient (bottom). Nuclear counterstain with DAPI or hematoxylin. **F.** pS6 and GFAP identifies giant cells in 230 days-old organoids comparable to giant cells in patient tubers. Giant cells in organoids express Vimentin, as shown in patients. Note that giant cells can be distinguished from tumors by their lower expression of Ki67. Nuclear counterstain with DAPI or hematoxylin. **G.** Tumors identified as pS6- and Ki67-positive areas are found in Patient 1 (N (batches) =5, n (organoids) =11, mean=1.9, SD=1.08) and Patient 2 (N=4, n=11, mean=5.5, SD=2.7) *TSC2*^+/−^-derived organoids at 110-days. Control organoids of Patient 1 (N=4, n=11) and two clones of repaired Patient 2 (clone 1 N=3, n=11; clone 2 N=2, n=8) showed no tumors. **H.** Percentage of tumor area of the total organoid area reveals similar tumor burden for organoids derived from both *TSC2*^+/−^ Patients (Pat.1 mean=15.7%, SD=14.2%; Pat.2 mean=17.1%, SD=11.3%; p=0.82, t-test). **I.** Area of the soma in Map2+ neurons shows that dysmorphic-appearing neurons in *TSC2*^+/−^-derived organoids are roughly 4-fold larger then Map2+ neurons in control organoids (Pat.1 Ctrl N=2, n=4, 858 neurons, mean= 76.9, SD=34.8; Pat.1 *TSC2*^+/−^ N=6, n=11, 241 dysmorphic neurons, mean=354.5, SD=180.4, Pat.1 Ctrl vs. *TSC2*^+/−^ p<0.0001; Pat.2 Ctrl1 N=2, n=4, 433 neurons, mean=81.4, SD=38; Pat.2 Ctrl2 N=2, n=5, 597 neurons, mean=70.51, SD=23.8; Pat.2 *TSC2*^+/−^ N=2, n=6, 240 dysmorphic neurons, mean=382.7, SD=159.8, Pat.2 Ctrl vs. *TSC2*^+/−^ and Pat.2 Ctrl vs. *TSC2*^+/−^ both p<0.0001; test: Ordinary One-Way ANOVA). **J.** Cell area of pS6-positive cells shows significantly enlarged pS6 cells in *TSC2*^+/−^-derived organoids, as compared to cells in either isogenic control. (Pat.1 Ctrl N=3, n=6, 1081 cells, Mean rank=1341; Pat.1 *TSC2*^+/−^ N=2, n=7, 810 cells, Mean rank=2274; Pat.2 Ctrl1 N=1, n=4, 608 cells, Mean rank=1291; Pat.2 *TSC2*^+/−^ N=3, n=12, 1146 cells, Mean rank=2241; Pat.1 Ctrl vs. *TSC2*^+/−^ p<0.0001, Pat.2 Ctrl vs. *TSC2*^+/−^ p<0.0001, Pat.1 Ctrl vs. Pat.2 Ctrl p>0.9999, Pat.1 *TSC2*^+/−^ vs. Pat.2 *TSC2*^+/−^ p>0.9999; Kruskal-Wallis test with Dunn’s multiple comparisons test) (Scale bars: C and D: 500μm; E: 20μm; F: 50μm)

Consistent with previous results^18^, we found no obvious differences between genotypes within the first 90 days of culture corresponding to early phases of neurodevelopment (Ext. Data Fig. 2). 110 days after embryoid body (EB) formation, however, nodular aggregates of cells expressing the proliferative marker Ki67 and the mTOR activation marker phospho-S6 (pS6) formed in *TSC2*^+/−^ organoids cultured in H-medium (Fig. 1D, G, H and Ext. Data Fig. 2). These structures morphologically resembled SENs or SEGAs that grow along the lateral ventricle in TSC patient brains^25–27^ (Fig. 1C). SEN/SEGAs are neurogliomal tumors that have been proposed to originate from an unknown population of neural stem cells (NSCs)^28^. To probe for an NSC origin of SEN-like tumors in organoids we stained for the classical NSC markers Nestin and Ascl1. Both proteins were expressed in tumors (Ext. Data Fig. 7A) indicating a neural progenitor identity of SEN/SEGAs.

To determine whether we can also recapitulate pathological cell types of cortical tubers, we analyzed organoids cultured for 120 to 150 days in L-medium. In organoids derived from *TSC2*^+/−^ cells, we found neurons with an enlarged soma and thickened processes, similar to dysmorphic neurons in cortical tubers (Fig. 1E, I). These neurons stained for the neural marker MAP2 and were not present in control organoids. After prolonged maturation in L-medium for about 230-days, clusters of enlarged pS6 positive cells appeared (Fig. 1F and Ext. Data Fig. 3). The morphology as well as the expression of markers like GFAP and Vimentin was reminiscent of giant cells (GCs), the second main pathological cell type of cortical tubers^7,9^, which had characteristically low proliferation rates (Fig. 1F). Thus, organoids derived from *TSC2*^+/−^ hiPSCs recapitulate the major histopathological features found in the brain of TSC patients.

## Single cell analysis of *TSC2*^+/−^ organoids reveals a caudal progenitor population

Since our organoid model recapitulated major disease features, we sought to identify the elusive common cell-of-origin for both cortical tubers and subependymal tumors^21^. We performed single cell RNA sequencing (scRNA-Seq) of 110 days-old organoids derived from *TSC2*^+/−^ and isogenic Ctrl (*TSC2^+/+^*) cells from Patient 1 cultured in L-medium. Unsupervised clustering overlaid on UMAP projection distinguished major cell types previously described in cerebral organoids^2,29^ (Fig. 2A and B, Ext. Data Fig. 4A).

**Fig. 2.**
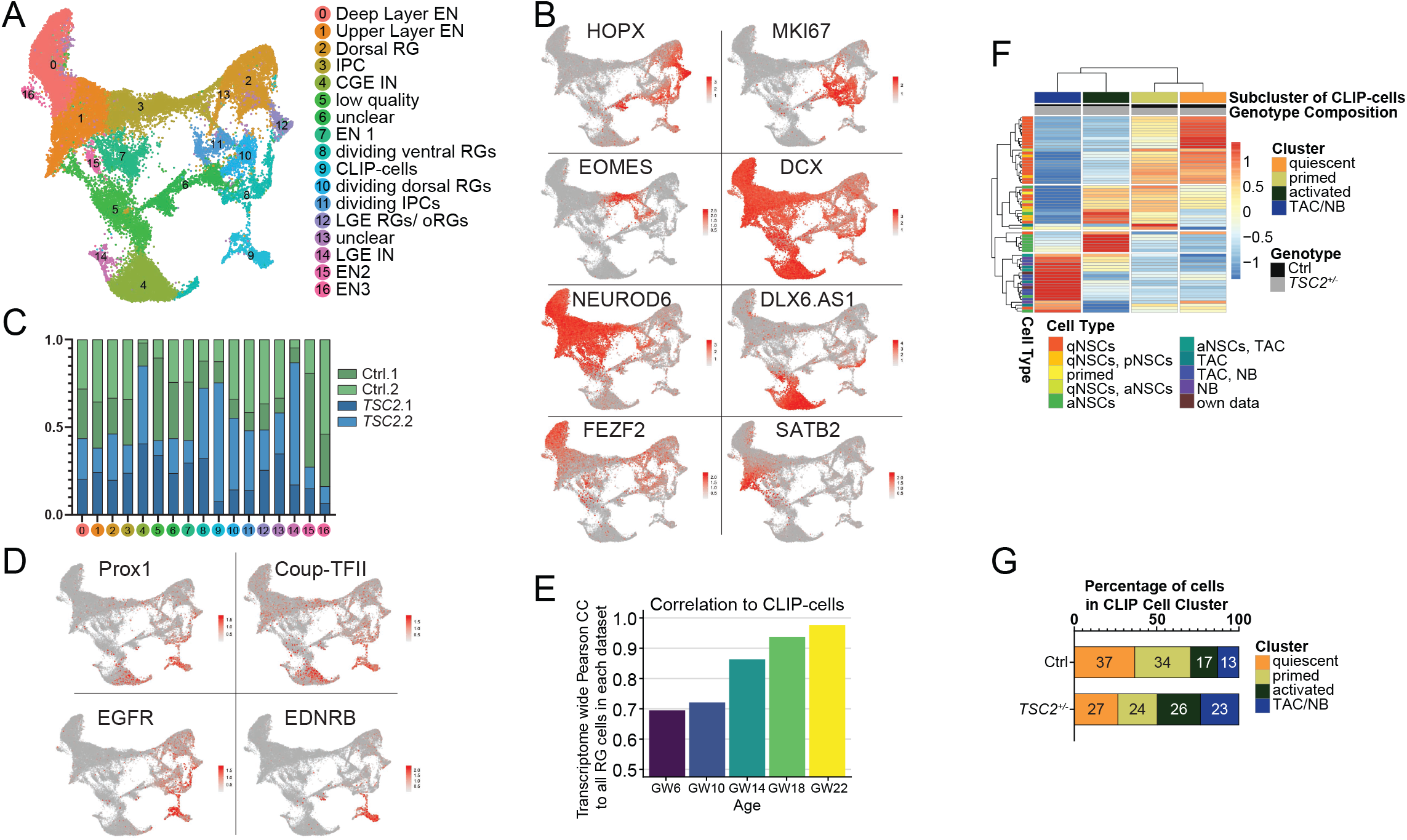
scRNA-seq identifies caudal late interneuron progenitor expanded in *TSC2*^+/−^ organoids. **A.** UMAP projection of all organoid cells from *TSC2*^+/−^ and *TSC2*^+/+^ (Ctrl) genotypes separates known cell types and a cluster of previously not identified progenitors, the CLIP cells (cluster 9). See Ext. Data Fig. 4A for expression of characteristic markers from differential gene expression analysis that allowed the identification of each cluster. **B.** Expression of classical cell type-specific markers: HOPX (outer radial glia, oRG), MKI67 (dividing cells), EOMES (also TBR2, dorsal intermediate progenitors), DCX (pan-neuronal), NEUROD6 (excitatory neurons, EN), DLX6.AS1 (inhibitory neurons), FEZF2 (deep layer EN), and SATB2 (upper layer EN). **C.** The cellular composition of each cluster from *TSC2*^+/−^ (Blue) and Ctrl (Green) genotypes shows that specific cell types are expanded in *TSC2*^+/−^ organoids. The shades of green and blue show cells from individual organoids. All cell numbers were down-sampled to the smallest dataset. The expanded cell types included interneurons (cluster 4, 14), as well as CLIP-cells (cluster 9) and dividing ventral RGs (cluster 8). Note that LGE RGs/ oRGs (cluster 12) showed similar cell numbers within each genotype. **D.** The expression of CGE-associated markers Prox1, Coup-TFII and EGFR in the CLIP-cells indicate their CGE origin. CLIP-cells also show other unique markers, such as Endothelin Receptor B (EDNRB). See Ext. Data Fig. 4B for expression of more markers. **E.** Transcriptome wide Pearson Correlation of organoid derived CLIP-cells to fetal RGs of different gestational ages derived from primary dataset29. Organoid CLIP-cells show the highest similarity to the oldest dataset (GW22). **F.** Clustering of all CLIP-cells and a genes discriminating different activation states of progenitors (genes derived from published scRNA-seq studies). For simplification, the average expression of each gene per group of cells is shown (See Ext. Data Fig. 5A and B for detailed heatmap). Genes were annotated from quiescent over primed and activated NSCs to transient amplifying cells (TAC) and neuroblasts with shared genes on each transition as in The heatmap identified four gene signatures (on the left side, from top to bottom): 1. quiescent, 2. mixed quiescent, primed and activated, 3. activated and 4. TAC and neuroblast genes. Based on distinct expression of these genes, we identified four cell sub-clusters, from left to right: TAC/ NBs, activated-, primed-, and quiescent NSCs. The percentage of each genotype contributing to the respective cell group is shown in gray/black on top of the heatmap. **G.** Percentage of each activation state within CLIP-cells from Ctrl and *TSC2*^+/−^ organoids. In Ctrl, most cells are quiescent or primed (q=37%, p=34%, a=17%, TAC/NB=13%), while in *TSC2*^+/−^ organoids CLIP-cells are biased towards activated states (q=27%, p=24%, a=26%, TAC/NB=23%).

Next, we identified cell types strongly expanded in *TSC2*^+/−^ organoids compared to Ctrl. We found that interneurons (cluster 4 and 14) and dividing cells of ventral origin (cluster 8) showed vastly increased contribution from *TSC2*^+/−^ organoids. Among progenitors, lateral ganglionic eminence (LGE)-derived radial glia (RGs) (cluster 12) were unchanged (Fig. 2A and C, Ext. Data Fig. 4A), but a previously undescribed cluster of progenitor cells was strongly enriched in *TSC2*^+/−^ cells (cluster 9, Fig. 2C, Ext. Data Fig. 4B). This cluster was characterized by the expression of genes associated with the caudal ganglionic eminence (CGE) like *COUP-TFII*, *PROX1* and the EGF receptor (*EGFR*, Fig. 2C). These cells also expressed a unique set of genes (Ext. Data Fig. 4B) such as the Endothelin Receptor B (*EDNRB*, Fig. 2C). To analyze whether this neural progenitor is also found *in vivo* we compared our data to a published scRNA-Seq dataset of the developing human fetal brain^30^. After integrating these data with our dataset, unsupervised clustering identified the same cluster in fetal brains (cluster 18 Ext. Data Fig. 4D, E). Correspondingly, we identified EGFR positive cells in immunostainings of the CGE at 21 gestational weeks (GW) (Ext. Data Fig. 5).

An interesting feature of both cortical tubers and tumors is the late developmental onset, both forming around mid-gestation^25^. As this is recapitulated in our organoid model, we analyzed whether the newly identified neural progenitor cells (NPCs) in organoids (cluster 9, Fig. 2A) were corresponding to fetal progenitors of a particular age. Therefore, we compared the transcriptome of fetal and organoid NPCs. We found the highest correlation of the newly identified caudal NPCs to RGs originating from late mid-gestational ages (Fig. 2D). Furthermore, co-clustering revealed that most fetal cells within the corresponding cluster (cluster 18, Ext. Data Fig. 4E) originated from gestational week (GW) 22 (Ext. Data Fig. 4F) indicating the late signature of this cell type. Based on these characteristics, the newly identified cells were named caudal late interneuron progenitors or CLIP-cells.

## CLIP-cells are activated in *TSC2*^+/−^

mTOR signaling regulates quiescence and activation in a variety of stem cell systems^31–33^. In order to understand whether altered mTOR signaling could affect the balance between quiescence and activation in *TSC2*^+/−^ CLIP-cells, we used multiple scRNA-Seq studies to derive a gene list that discriminates between activated and quiescent states (Ext. Data Table 2)^34–37^. We performed hierarchical clustering of all identified CLIP-cells and the expression of genes on this list (Fig. 2E and Ext. Data Fig. 5A, B). This defined four sub-clusters of quiescent-, primed-, activated- and transient amplifying cells (TAC) / neuroblasts (Fig. 2E, Ext. Data Fig. 5A, B). In *TSC2*^+/−^ organoids, activated sub-populations were increased while quiescent cells were reduced (Fig. 2E, Ext. Data Fig. 5A, B and C). Our analysis identified EGFR expression as a marker for activated CLIP-cells. Immunostainings revealed more EGFR positive cells in *TSC2*^+/−^ organoids, confirming the scRNA Seq results (Ext. Data Fig. 5D). This suggests a role for the mTOR pathway in controlling CLIP cell activation that could explain why *TSC2*^+/−^ organoids show an increase in CLIP cells and in CGE-derived interneurons (Fig. 2B).

## TSC tumors express CLIP cell markers

In order to investigate whether CLIP-cells are the neural progenitors causing neurogliomal TSC tumors, we stained *TSC2*^+/−^ organoids for the CLIP-cell markers identified in our scRNA-seq dataset (Ext. Data Fig. 4B). Among these are the CGE-associated markers COUP-TFII, PROX1, SCGN and EGFR, as well as the unique markers EDNRB and prostaglandin D synthase (PTGDS).

To test for a CGE-origin we investigated expression of EGFR and COUP-TFII in tumors. EGFR (Fig. 3A) was co-expressed with COUP-TFII (Ext. Data Fig. 7C and D) in SEN-like tumors within *TSC2*^+/−^ organoids. Consistent with previous observations^38^, we also detected EGFR expression in surgically removed SEGAs (Fig. 3A). To assess unique CLIP-cell markers, we stained organoids for EDNRB and PTGDS, in addition to the CGE marker PROX1. EDNRB (Fig. 3B) was both co-expressed with PROX1 (Ext. Data Fig. 8A) in organoid tumors and detected in SEGAs resected from patient brains (Fig. 3B). Similarly, PTGDS was expressed together with COUP-TFII and PROX1 in *TSC2*^+/−^-derived organoid tumors (Ext. Data Fig. 9B) and in CLIP-cells in Ctrl organoids of the same age (Ext. Data Fig. 9A). Finally, we tested expression of SCGN, a marker for the CGE derived interneurons. SCGN was expressed in tumors in organoids (Ext. Data Fig. 10D) and in resected SEGA samples (Ext. Data Fig. 10E), indicating that CLIP-cells in tumors can still produce interneurons.

**Fig. 3.**
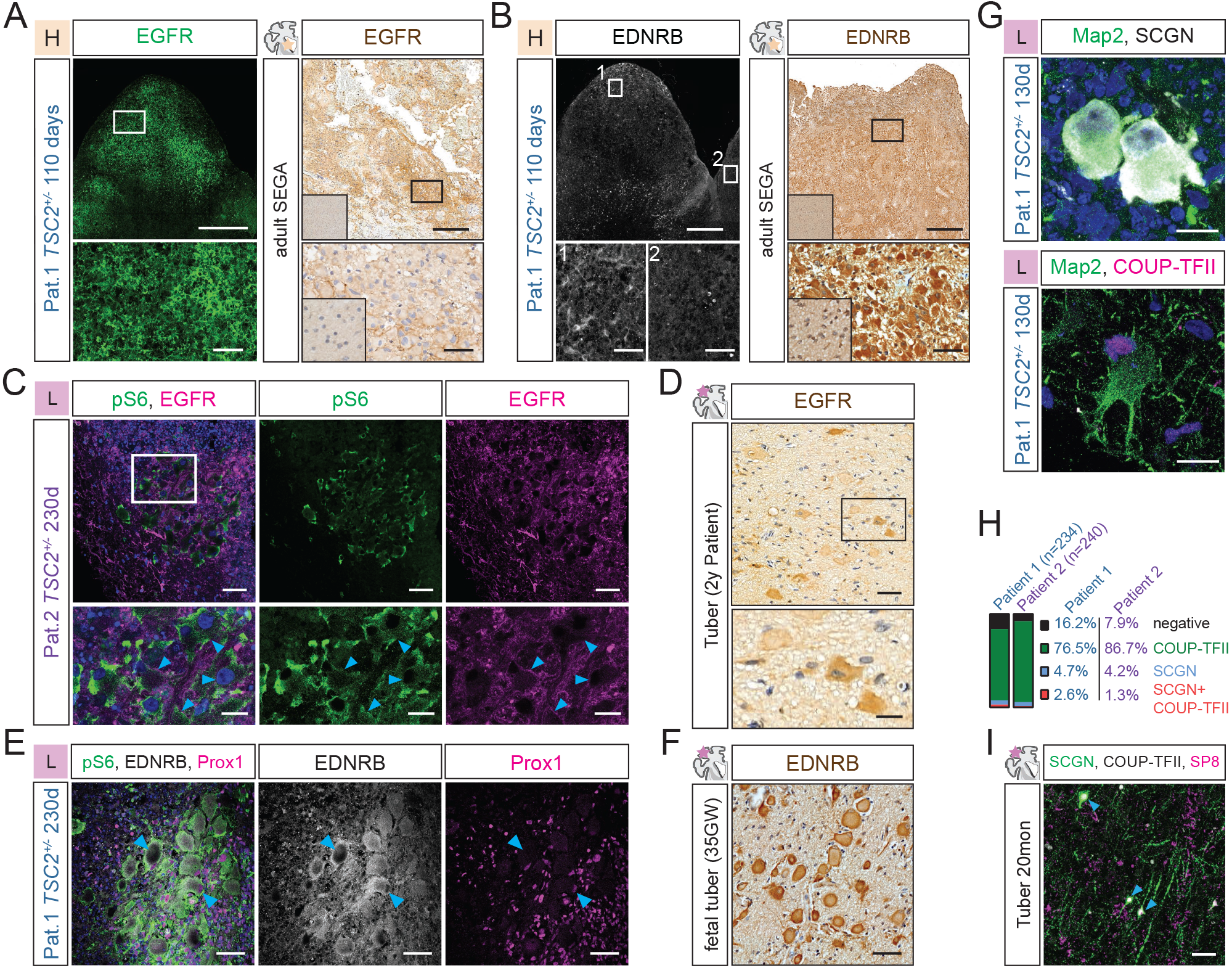
CLIP-cells generate all abnormal cell types of TSC brain pathology. **A.** Left: Immunostaining for EGFR identifies tumors in 110 days-old *TSC2*^+/−^-derived organoids grown in H-medium. See Ext. Data Fig. 7C and D for co-staining with COUP-TFII. Lower panel is higher magnification of tumor region. Right: EGFR expression is present in adult SEGAs. Inset in primary tissue staining shows that a cortex from an healthy control is negative for EGFR. Lower panel is higher magnification of tumor region. **B.** Left: EDNRB is expressed in tumors in 110 days-old *TSC2*^+/−^-derived organoids. Zoom-in #1 shows a tumor region expressing EDNRB and #2 shows a negative region outside of the tumor. See Ext. Data Fig. 8A for co-staining with PROX1 and pS6. Right: EDNRB expression is also found in adult SEGAs. Lower panel is higher magnification of tumor region. Inset in primary tissue staining shows healthy control cortex negative for EDNRB. **C.** Giant cells in 230 days-old *TSC2*^+/−^-derived organoids grown in L-medium co-express pS6 and EGFR. Note that EGFR is localized to the membrane of giant cells. Blue arrows mark examples of EGFR-positive membrane. Lower panel shows higher magnification **D.** EGFR is found in giant cells in a resected postnatal cortical tuber of a 2 years-old patient. **E.** pS6, EDNRB and PROX1 are co-expressed in Giant cells in in 230 days-old organoids *TSC2*^+/−^-derived organoids grown in L-medium. **F.** EDNRB is also expressed in giant cells of a fetal TSC tuber. See Ext. Data Fig. 8E for postnatal tuber example. **G.** Dysmorphic neurons in *TSC2*^+/−^-derived organoids in L-medium express SCGN and/or COUP-TFII and show enlarged soma and thickened processes. See Ext. Data Fig. 10A for Ctrl and further examples. **H.** Quantification of dysmorphic neurons expressing CGE-associated markers in *TSC2*^+/−^-derived organoids. Note that dysmorphic neurons are only found in *TSC2*^+/−^, but not in controls (Fig. 1I). **I.** SCGN, COUP-TFII and SP8 in a resected cortical tuber of a 20 months-old TSC case identifies enlarged, dysmorphic neurons (blue arrows). See Ext. Data Fig. 10B for healthy control. See Ext. Data Fig. 10C for fetal tuber example. (Scale bars: overview images A, B: 500μm; Zoom-in A, B.: 50μm; overview C, D, E, F, I: 50μm; Inset C, D: 20μm; G and H: 15μm)

Taken together these data suggest that CLIP-cells might be the developmental cell-of-origin of TSC tumors and constitute the neurogliomal identity of SENs and SEGAs together with their interneuron progeny.

## Giant cells in cortical tubers express CLIP-cell markers

To address whether CLIP-cells also generate the giant cells found in cortical tubers, we tested the same markers in tuber-like structures in organoids and in patient derived brain tissue. In 230 days-old *TSC2*^+/−^-derived organoids cultured in L-medium almost all pS6-positive giant cells stain positive for EGFR (Fig. 3C). Correspondingly, EGFR was also expressed in giant cells within postnatally resected tubers (Fig. 3D). To address expression of CLIP-cell specific markers in giant cells, we stained for EDNRB and PTGDS as well as the CGE-associated markers PROX1 and COUP-TFII. We found co-expression of EDNRB and PROX1 (Fig. 3E) in giant cells in organoids and high expression of EDNRB in giant cells in fetal tubers (Fig. 3F) as well as in postnatally resected tubers (Ext. Data Fig. 8E). Likewise, we detected PTGDS together with PROX1 and COUP-TFII (Ext. Data Fig. 9C) in giant cells in *TSC2*^+/−^-derived organoids. Thus, giant cells in organoids and tubers express CLIP-cell specific proteins, suggesting that CLIP-cells also give rise to giant cells characteristic of cortical tubers.

## Dysmorphic neurons found in cortical tubers are CGE-derived interneurons

To investigate whether dysmorphic neurons found in cortical tubers are related to CLIP-cells, we stained for the CGE markers SCGN and COUP-TFII. Most dysmorphic neurons (identified by MAP2 staining, see Fig. 1E) expressed COUP-TFII and/or SCGN (Pat.1: 84%, Pat.2 92%; Fig. 3G and H, Ext. Data Fig. 10A). Accordingly, we found expression of SCGN in fetal cortical tubers (Ext. Data Fig. 10C) and postnatally resected tubers (Fig. 3I and Ext. Data Fig. 10B). SCGN and COUP-TFII positive neurons in tubers differed strikingly from age-matched healthy control cases, showing thickened dysmorphic processes and enlarged soma (Fig. 3J and Ext. Data Fig. 10B). These data demonstrate that dysmorphic neurons in organoids and in TSC patients are CGE-derived interneurons, suggesting that they originate from CLIP-cells.

Taken together, we propose that TSC arises from cell-type specific dysregulation of neurogenesis in the CGE and that CLIP-cells are the common cell-of-origin of both the pathological cell types found in tumors and those seen in cortical tubers.

## Bi-allelic *TSC2* inactivation is dispensable for formation of TSC lesions

As pathological cell types form reproducibly in *TSC2*^+/−^ organoids, it is unlikely that their generation requires loss of the second *TSC2* allele. Bi-allelic inactivation almost exclusively occurs by copy-neutral loss-of-heterozygosity (cnLOH), and it has been proposed to cause SEN/SEGAs in TSC patient brains^21,22^. Cortical tuber lesions, in contrast, only very rarely harbor second-hit events^19–21,39^. To assess the role of second-hit mutations in the TSC organoid model, we first investigated regions with giant cells resembling cortical tubers. In 230 days-old *TSC2*^+/−^ organoids, TSC2 protein was detected in over 98% of the giant cells using an antibody recognizing only the wild type TSC2 variant (Patient 1 98.4%, Patient 2 98.7%; Ext. Data Fig. 11A-C). Similarly, and consistent with previous data^40,41^, TSC2 protein was expressed in cortical tubers resected from TSC patients (Ext. Data Fig. 11D). This suggests that second-hit events are not a prerequisite for tuber formation.

To determine whether bi-allelic inactivation is also dispensable for the initiation of tumor lesions, we directly tested the mutational status of 135 to 160-day old *TSC2*^+/−^-derived organoids. For this, we enriched activated CLIP-cells from both Ctrl and tumor containing mutant organoids (Fig. 4A) by Fluorescent activated cell sorting (FACS) for EGFR expression. Both total cell number, as well as absolute and relative abundance of EGFR-positive cells were increased in *TSC2*^+/−^-derived organoids (Ext. Data Fig 12). DNA sequencing across the genomic region carrying the *TSC2* mutation showed that the EGFR-positive cells in organoids derived from *TSC2*^+/−^ cells remained heterozygous in the majority (7/11) of cases (Fig. 4B and C and Ext. Data Fig. 13A). Only in one of the tumors, we identified a complete cnLOH while three tumors showed a partial cnLOH (Fig. 4C). To further analyze recombination events in homozygous TSC tumors, we performed whole genome sequencing (WGS) on two tumor samples with cnLOH. WGS confirmed cnLOH and revealed that large regions of chromosome 16 around the *TSC2* locus had become homozygous by recombination (Fig. 4D and E and Ext. Data Fig.12B). No other mutations were observed in the cnLOH and the general mutation frequency was low, as previously reported in TSC tumors^21^. Notably, tumors that had become homozygous have more EGFR-positive cells (Ext. Data Fig 13D), indicating they are larger. To further analyze, if cnLOH indeed occurred preferentially in larger tumors, we performed Laser-Capture-Microdissection (LCM) of small and enlarged tumors. While small tumors remained heterozygous, large tumors exhibited cnLOH events (Ext. Data Fig. 13F). These data suggest that a second-hit at the *TSC2* locus is not required for tumor initiation but occurs at later stages of tumor progression.

**Fig. 4.**
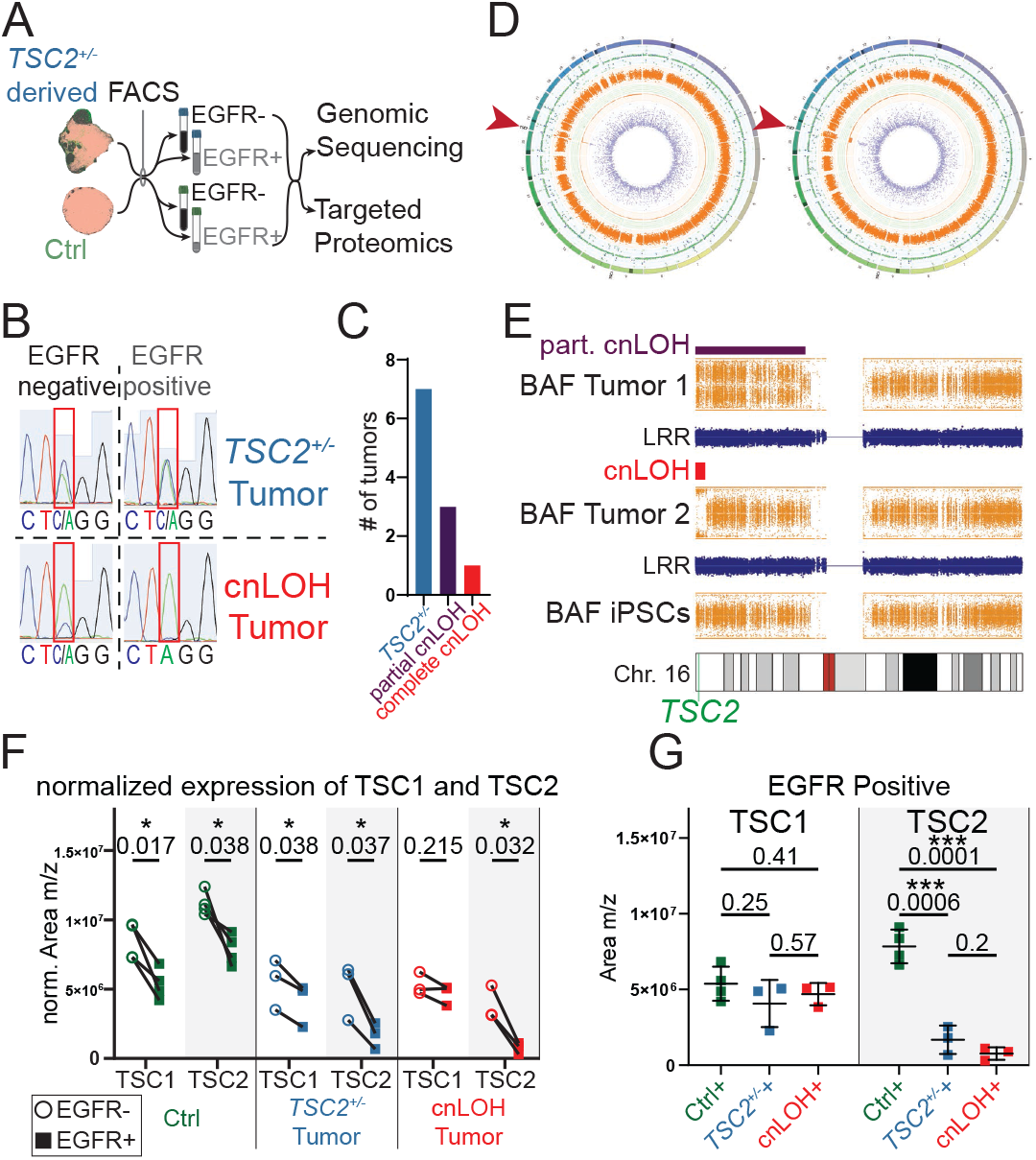
Sequencing and targeted PRM on TSC tumors. **A.** FACS strategy enriching for tumor population in *TSC2*^+/−^-derived organoids using EGFR expression. **B.** Mutation locus in the TSC2 gene from matched EGFR-negative and -positive samples of heterozygous and cnLOH tumor show loss of the WT (C) allele. **C.** 7 of 11 TSC2+/--derived organoids genotyped after FACS were heterozygous, three showed partial cnLOH, and only one showed complete cnLOH, reflecting different tumor stages. **D.** Circos plots of WGS of two EGFR-positive tumor samples from *TSC2*^+/−^-derived organoids with partial and complete cnLOH samples using *TSC2*^+/−^-iPSCs as a reference. There are no alterations on other chromosomal loci except for TSC2 locus on Chr. 16 (red arrows) compared to iPSCs. See panel E for close-up of Chr. 16 and Ext. Data Fig. 13 for further explanation. **E.** More detailed depiction of cnLOH around TSC2 locus. BAF (orange) indicates different regions of loss-of-het-erozygozity around TSC2 locus along chromosome 16 for both tumors while iPSC reference remains heterozygous. Unchanged log-R ratio (LRR, blue) indicates that no aneuploidy occurred, thus copy numbers remained the same. This gives the definite diagnosis of a copy-neutral loss-of-heterozygosity, characteristic for TSC tumors. Note that in tumor 2 all cells have cnLOH shown as a complete shift of BAF along the Y-axis. In tumor 1 the shift is less clear indicating mixed cnLOH and *TSC2*^+/−^ tumor cells. **F.** Normalized protein expression of TSC1 and TSC2 in matched EGFR-negative (circle) and EGFR-positive (square) populations sorted from 3 different genotypes (Ctrl; *TSC2*^+/−^ tumor; cnLOH tumor). Protein quantification shows that EGFR-positive cells have decreased TSC2 expression in all cases, while TSC1 expression is less extreme. The loss of TSC2 protein in EGFR+ samples is further decreased by the stepwise loss of the healthy TSC2 allele (*TSC2+/+* in Ctrl, *TSC2*^+/−^, and *TSC2*/−). (ctrl N=2, n=4; *TSC2*^+/−^ N=2, n=3; cnLOH N=2, n=3; matched EGFR-positive and -negative for each sample; p-values of paired t-test above samples). **G.** Directly comparing TSC1 or TSC2 protein levels across genotypes reveals that EGFR-positive tumor samples drastically downregulate TSC2, more than expected by the 50% loss in gene dosage. TSC1 levels are similar in EGFR-positive populations of all three genotypes. Statistical Test: F: Paired t-test, G: unpaired t-test

## Activated CLIP-cells are uniquely susceptible to a heterozygous *TSC2* state

As bi-allelic *TSC2* inactivation was dispensable for both tuber and tumor formation, we asked why CLIP-cells are particularly sensitive to heterozygous TSC states. We hypothesized that low endogenous TSC complex levels make CLIP-cells uniquely susceptible to any further reduction of TSC1 or TSC2. We therefore measured TSC1 and TSC2 protein levels by targeted parallel reaction monitoring (PRM) in previously genotyped samples of FACS sorted EGFR-positive and -negative cells (Fig. 4 F, G and Ext. Data Fig. 14). Both in Ctrl and *TSC2*^+/−^-derived organoids, TSC1 and TSC2 levels were lower in EGFR positive samples than in EGFR-negative samples (Fig. 4F). However, while in EGFR-negative cells, TSC1 levels were reduced in *TSC2*^+/−^-derived organoids compared to Ctrl (Ext. Data Fig. 14C), in the EGFR-positive population TSC1 levels were similar between Ctrl and *TSC2* mutants (Fig. 4G). TSC2, in contrast, was significantly more downregulated in EGFR-positive cells in the *TSC2* mutant compared to the Ctrl population (Fig. 4G). Thus, while in Ctrl organoids both components of the TSC complex were equally reduced in activated CLIP-cells, loss of one functional *TSC2* allele lead to disproportional reduction of TSC2 in EGFR-positive cells. To confirm whether reduction of TSC2 protein is specifically found within tumors, we stained for TSC2. We found a focal reduction of TSC2 in SEN-like tumors, while surrounding regions show TSC2 expression (Ext. Data Fig. 11E). These results demonstrate, that CLIP-cells have particularly low levels of TSC proteins making them uniquely sensitive to mutations in TSC genes.

## TSC tumors depend on EGFR signaling

Both CLIP-cells and the resulting tumors express EGFR. To address the functional implications of the EGFR pathway for CLIP-cell proliferation and tumor progression, we developed a drug-testing assay in our organoid model. Current pharmacological treatment of SEN/SEGAs in TSC is based on the mTOR Complex 1 inhibitor Everolimus^42–47^. Reports of variable responses and tumor re-growth after discontinuation necessitate life-long treatment and require alternative therapeutic strategies. We hypothesized that targeting activated CLIP-cells in TSC tumors through the EGFR pathway might reduce tumor burden and/or size. 110 days-old organoids grown in H-medium were treated over the course of 30 days with the mTOR inhibitor Everolimus, the EGFR receptor tyrosine kinase inhibitor (RTKI) Afatinib, or the same concentration of DMSO as a control (Ext. Data Fig 15A). Reductions in tumor burden or size were determined by measuring areas co-expressing pS6 and EGFR. Everolimus treatment almost completely abolished tumor appearance in 140 days-old organoids (Fig. 5A, B). After Afatinib treatment, both tumor load and mean tumor size were significantly reduced when compared to untreated organoids (Fig. 5A and B, Ext. Data Fig. 15B). These data strongly support a role of EGFR signaling in TSC tumor biology and suggest that targeting the EGFR pathway could be used as an alternative treatment for TSC brain lesions.

**Fig. 5.**
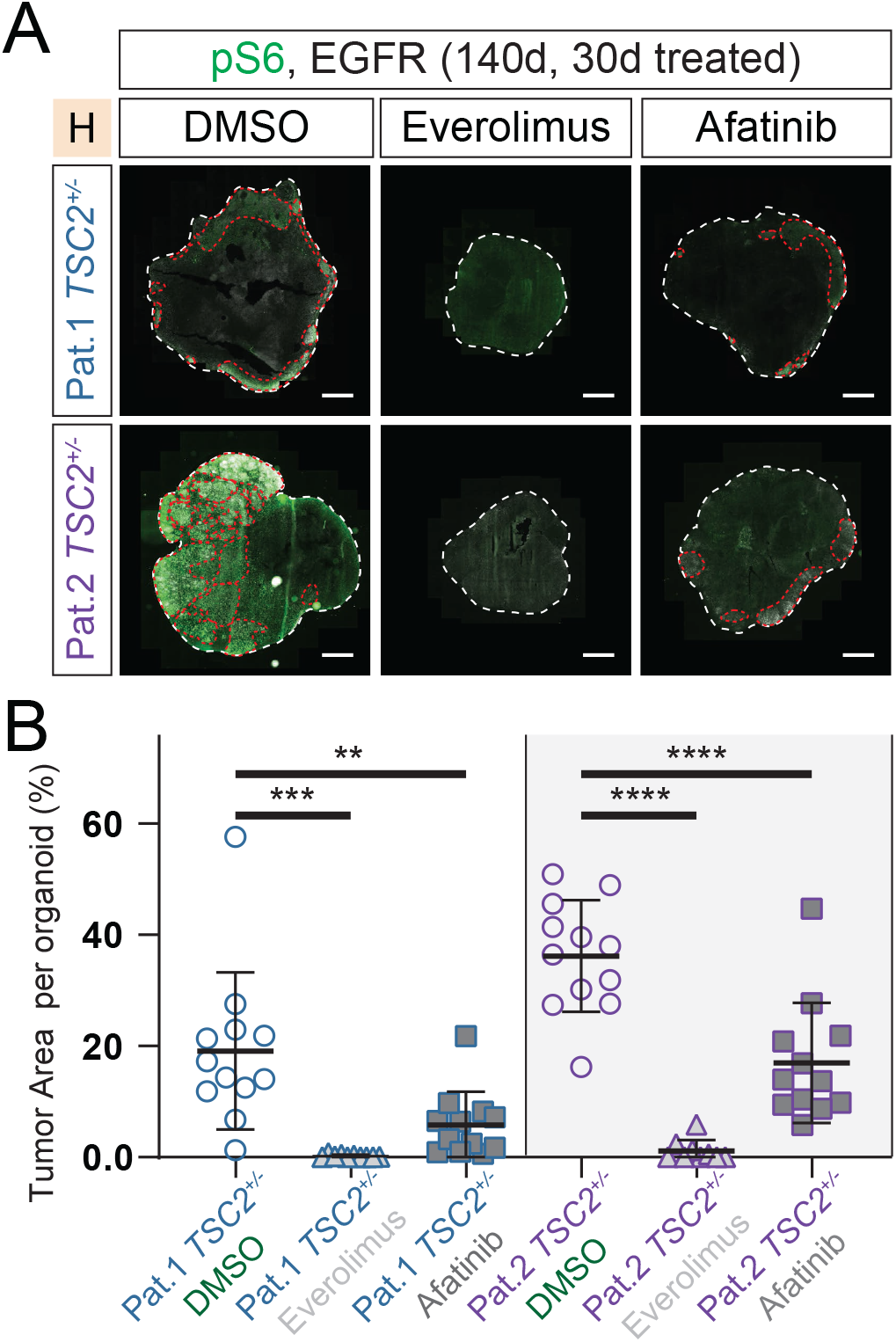
EGFR-inhibition reduces tumor burden. **A.** 30 days of treatment were initiated at 110 days after EB formation (see Ext. Data. Fig. 15A) Immunostaining for pS6 and EGFR identifies tumors (red lines) in the control group (DMSO) in both patients. Tumors are severely reduced by Afatinib treatment, and completely disappear after Everolimus treatment. **B.** Quantification of tumor area per organoid. All sections of each organoid stained on one slide were used for quantification. Tumor was identified as region of overlapping pS6 and EGFR staining. While tumors are detected in DMSO-control for both patients, Afatinib or Everolimus treatment both reduce tumor burden significantly. (N=3, n=12 for Pat.1 and Pat.2 *TSC2*^+/−^-DMSO and Afa; N=3, n=10 for Pat.1 *TSC2*^+/−^-Eve; N=3, n=9 for Pat.2 *TSC2*^+/−^-Eve; Pat.1 *TSC2*^+/−^-DMSO mean=19.1%, SD=13.5%; Pat.1 *TSC2*^+/−^-Eve mean=0.19%, SD=0.25%; Pat.1 *TSC2*^+/−^-Afa mean=5.81%, SD=5.69%; Pat.2 *TSC2*^+/−^-DMSO mean=36.19%, SD=9.62%; Pat.2 *TSC2*^+/−^-Eve mean=1.12%, SD=1.86%; Pat.2 *TSC2*^+/−^-Afa mean=16.99%, SD=10.36%; Pat.1 *TSC2*^+/−^-DMSO vs. Eve p=0.0001, Pat.1 *TSC2*^+/−^-DMSO vs. Afa p=0.0081, Pat.2 *TSC2*^+/−^-DMSO vs. Eve p<0.0001, Pat.2 *TSC2*^+/−^-DMSO vs. Afa p<0.0001; Ordinary one-way ANOVA, Tukey’s multiple comparisons test) (Scale bars: A.: 1mm)

## Discussion

While numerous studies in mouse models and a recent spheroid model have defined loss-of-heterozygosity as a requirement for TSC-like phenotypes^13–18^, our data suggest that bi-allelic inactivation is dispensable for disease initiation. Instead, cnLOH events occur during tumor progression and thus are secondary in tumor pathogenesis. Our observations are in line with reports in patients that identify bi-allelic inactivation in subependymal tumors but not cortical tubers in the brain^19-22,39,48^.

TSC phenotypes can first be detected around late mid-gestation^25^. In agreement with this, we identify CLIP-cells, a late progenitor cell that emerges only after the early phases of neurogenesis have been completed as the cell-of-origin for TSC. CLIP cells show unique susceptibility towards changes in mTOR signaling that cause TSC brain pathology. The identification of CLIP-cells as the common cell-of-origin for TSC brain lesions is the first demonstration of a single founder population giving rise to two genetically and morphologically distinct phenotypes during brain development. Thus, our findings explain both the transcriptional commonalities^21^ and the differences in mutational status between TSC lesions, resolving two long-standing questions in the TSC field.

Our work demonstrates that studying the mechanisms of neurodevelopmental diseases like tuberous sclerosis complex can give insights into human-specific processes in brain development. The specific expression profile of CLIP-cells namely a caudal and mid-gestational signature has not been described in the human brain before. However, some of the identified genes, like EDNRB, have been found in adult neural stem cells responsible for postnatal neurogenesis in other mammals like the mouse^35^. In mice, however, these cells are only permissive for TSC phenotypes in a *Tsc1*^null^ state^14^. Additionally, SCGN interneurons as produced by CLIP-cells do not exist in the mouse^49^. Thus, our data raise the exciting possibility, that the population uniquely susceptible to mutations in the TSC-mTOR pathway only exists in the human brain. This not only explains why heterozygous *Tsc1/Tsc2* mouse models do not develop TSC phenotypes, but also shows the requirement for human models in order to study neurodevelopmental disease.

The CGE has previously been described as a source of human specific changes in neurodevelopment: Firstly, CGE-derived interneurons contribute to the human brain in much higher percentages compared to rodents^50,51^. Secondly, late-migrating SCGN interneurons from the CGE have been observed in humans, but not in mice^49^. Thirdly, a stream of interneurons also containing CGE-derived interneurons, continues to migrate into the human brain even after birth^6^. However, the relevance of this population for human neurodevelopment and disease has remained elusive. We identify expression of CGE markers in dysmorphic neurons in *TSC2*^+/−^-derived organoids and TSC patients. Together with our characterization of CLIP-cells emerging around mid-gestation, this suggests that CLIP-cells could be the progenitor population giving rise to human specific late-migrating interneurons derived from the CGE.

The finding that CLIP-cells and their interneuron progeny are relevant for disease can help to understand the TSC associated specific neuropsychiatric symptoms. The range of neuropsychiatric impairments in TSC patients consolidated as tuberous sclerosis-associated neuropsychiatric disorders, or TAND comprises a variety of complex conditions involved in higher brain functions like autism or attentional deficits and other specific neuropsychological defects^52^. It will be interesting to further investigate the role of interneurons derived from CLIP-cells in the alteration of these human specific traits. Our work highlights the importance of studying late stages of the prolonged and complex human neurogenesis in order to better understand human neurodevelopmental disorders.

## Data Availability Statement

All sequencing data is submitted to the European genome-phenome archive (EGA) and will be made available under restricted access before publication.

## Code Availability Statement

All custom code used to for data analysis will be made available on GitHub at https://github.com/vertesy/TSC2-CLIP.

## Acknowledgements

We thank Paul Möseneder, Hilary Eleanor Gustafson and Simone Wolfinger for excellent technical assistance; the IMBA Stem Cell Core Facility and Chukwuma Allison Agu for generation of IPS Cell Lines; Bartlomiej Gebarski and Alex Vogt for library preparation and sequencing performed at the VBCF NGS Unit (https://www.viennabiocenter.org/facilities/next-generation-sequencing/); the Genome Engineering Unit of VBCF ProTech facility (https://www.viennabiocenter.org/facilities/protein-technologies/) for cloning of Crispr Guides, purified SpCas9 protein and assistance during isogenic control line preparation; Karel Stejskal and Elisabeth Roitinger for mass spectrometry and analysis performed in the IMBA/IMP Mass spectrometry facility; the IMBA/IMP Biooptics facility for valuable assistance with FACS sorting and Image acquisition; Adrian Mancebo Gimenez of the VBCF HistoPathology facility for immunohistochemistry on primary material; Arabella Meixner for coordinating the ethical approvals involved in this study; Robert Diehm; Gregor Kasprian for providing patient MRI; the KIN Biobank of the MUV for providing paraffin-embedded fetal tissues; Kurtis Auguste for providing primary material; Vicente Elorriaga Benavides for initial work on primary material and Oliver Wüseke for establishing initial contact with the MUV. We especially thank all patients and their families for participating in this study or donating tissue.

A.V. was supported by an EMBO Fellowship (ALTF-1112-2019). Work in J.A.K.’s laboratory is supported by the Austrian Federal Ministry of Education, Science and Research, the Austrian Academy of Sciences, the City of Vienna and the SFB F78 Stem Cell (F 7803-B). This project has received funding from the European Research Council (ERC) under the European Union’s Horizon 2020 research and innovation (695642).

## Author contributions

O.L.E, N.S.C., M.F. and J.A.K. designed the study and analysis. Experiments were performed by O.L.E., N.S.C., T.S., V.E.G., A.M.P., and J.C.. Data analysis was performed by O.L.E., N.S.C., A.V., M.N. and M.F.P.. The study was supervised by N.S.C. and J.A.K. The manuscript was prepared by O.L.E, N.S.C. and J.A.K. with input from all authors.

## Materials & Methods

### Patient sample selection

The study was approved by the local ethics committee of the Medical University of Vienna (MUV). Screened were all patients with TSC included into our TSC data registry. Study inclusion criteria were as follows: 1) TSC proven by clinical characteristics and molecular testing. 2) age between 0 and 18 years. 3) TSC-associated drug-resistant epilepsy. 4) continuous follow-up at the department of pediatrics TSC center for at least 2 years. After informed consent, 10 ml blood was collected from three selected patients for iPSC reprogramming. All clinical data were derived from the MUV TSC patient registry (e.g. gender, age, seizure frequency, seizure onset, pre- and postnatal MRIs and AEDs). Re-evaluation of formalin-fixed and paraffin-embedded (FFPE) TSC brain material (cortical tubers and SEGAs) of one included iPSC patient and several other TSC patients was performed. In addition, brain material of fetal autopsy cases with TSC were also included for comparative studies. Age- and region-matched autopsy tissue samples from patients without a history of any neurological disease including epilepsy served as control group.

### Generation of IPS cells

IPS cells were generated from PBMCs isolated from patient blood samples as described^1^.

Briefly, 10 ml blood was collected in sodium citrate collection tubes. PBMCs were isolated via a Ficoll-Paque density gradient and erythroblasts were expanded for 9 days. Erythroblast-enriched populations were infected with Sendai Vectors expressing human OCT3/4, SOX2, KLF4 and cMYC (CytoTune, Life Technologies, A1377801). Three days after infection, cells were switched to mouse embryonic fibroblast feeder layers and medium was changed to IPSC media (KoSR + FGF2) 5 days post-infection. 10 to 21 days after infection, the transduced cells began to form colonies that exhibited IPSC morphology. IPSC colonies were picked and passaged every 5 to 7 days. IPSCs were passaged 15 times before being transferred to the Cellartis DEF-CS 500 culture system (Takara).

### IPS cell culture

iPSCs cell culture was performed using the Cellartis DEF-CS 500 culture system (Takara) according to the manufacturers’ instructions. Briefly, cells were split every third day and 400.000 cells were seeded per well of a 6 well plate. Cells were banked at different passages. For experiments, cells were used from passage 40 to 50. Genomic integrity was analyzed at passage 40 on an Infinium PsychArray v1.3 (Illumina) and compared to data from PBMCs.

### Generation of isogenic control cell lines

Reprogramming of PBMCs from Patient 1 generated IPSC clones with the genotypes *TSC2*^+/−^ and *TSC2*^+/+^, indicating, that Patient 1 carried a mosaic heterozygous mutation in *TSC2*. Mosaicism was confirmed in PBMCs.

Isogenic control cell lines of patient 2 were generated using Crispr/Cas9. *S.pyogenes* Cas9 protein with two nuclear localization signals was purified as previously described^2^. gRNA transcription was performed with HiScribe T7 High Yield RNA Synthesis Kit (NEB) according to the manufacturer’s protocol and gRNAs were purified via Phenol:Chloroform:Isoamyl alcohol (25:24:1; Applichem) extraction followed by ethanol precipitation. The HDR template (custom ssODN (Integrated DNA Technologies)) was designed to span 100 bp up and downstream of the mutation site. For generation of isogenic control cell lines, IPSCs were grown in DEF-CS. Cells were washed with D-PBS−/−. TrypLE Select (Thermo Fisher Scientific) was added to cells and cells were incubated for 5 min at 37°C. Plate was gently tapped to dissociate cells, resuspended in supplemented DEF-CS medium and counted. 1.5 × 10^6^ cells were spun down and washed with D-PBS−/− once. Cells were resuspended in Buffer R of the Neon Transfection System (Thermo Fisher Scientific) at a concentration of 2×10^7^ cells/ml. 0.45 μl of sgRNA (1 μg/μl), 0.75 μl electroporation ready Cas9 protein (3μg/μl) and 6.3 ul of resuspension buffer were combined to for the Cas9/sgRNA RNP complex, reaction was mixed and incubated at 37°C for 5 min. 2 μl of the HDR template (100 μM) was added to the Cas9/sgRNA RNP complex and combined with the cell suspension. Electroporations were performed using a Neon^®^ Transfection System (Thermo Fisher Scientific) with 100 μl Neon^®^ Pipette Tips using the ES cells electroporation protocol (1400 V, 10 ms, 3 pulses). Cells were seeded in one well of a 6 well plate in supplemented DEF-CS (Takara). After a recovery period of 5 days, cells were FACS sorted in 96 well plates with 1 cell/ well. 48 h after sorting, 100 μl of fresh supplemented DEF-CS medium was added. After expansion, gDNA was extracted using DNA QuickExtract Solution (Lucigen) followed by PCR and Sequencing to determine efficient rescue.

### Organoid generation

To generate organoids from DEF-CS cultured iPSCs, 400.000 cells were seeded per well of a 6 well plate. After 2 days, cells were washed with DPBS−/− and incubated with 300 μl TryLE Express (Thermo Fisher Scientific) for 5 minutes. 1 ml supplemented Def-CS (basal medium + GF1,2,3) was added and iPSCs were triturated and subsequently transferred to a tube with 700 ul supplemented Def-CS. Cells were counted and the required number of cells was added to a new tube. Cells were spun down (120 × g) for 3 min and resuspended in the required amount of mTeSR (Stem Cell Technologies) + Rock Inhibitor (RI): 9000 cells were seeded per well of a low attachment 96 well plate in 150 μl mTeSR + RI. The resulting embryoid bodies were fed 2 and 4 days after EB generation with mTeSR without RI. At day 5 after EB generation, medium was replaced with Neural Induction medium consisting of DMEM/F12 (Thermo Fisher Scientific) with 1% N2 Supplement (Thermo Fisher Scientific), 1% MEM-NEAA (Sigma Aldrich), 1% Glutamax (Thermo Fisher Scientific) and 5 ug/ml Heparin. Medium was changed at day 7 and day 9 after EB generation. 10 days after EB generation, EBs were embedded in Matrigel (Corning) as described ^3^. Embedded EBs were cultured in High Nutrient Medium - Vitamin A (HN-A) consisting of 50% DMEM/F12 (Thermo Fisher Scientific), 50% Neurobasal (Thermo Fisher Scientific), 1 % N2 Supplement (Thermo Fisher Scientific), 2% B27 - Vitamin A (Thermo Fisher Scientific), 1% Glutamax (Thermo Fisher Scientific), 0,5% MEM-NEAA (Sigma Aldrich,), 1 % Penicillin/Streptomycin (Sigma Aldrich) and 0.025% Insulin solution (Sigma Aldrich). Organoids were cultured in stationary culture for 3 days. 13 days after organoid generation, medium was changed to High Nutrient Medium + Vitamin A (HN+A) consisting of 50% DMEM/F12 (Thermo Fisher Scientific), 50% Neurobasal (Thermo Fisher Scientific), 1 % N2 Supplement (Thermo Fisher Scientific), 2% B27 + Vitamin A (50X, Thermo Fisher Scientific), 1% Glutamax (Thermo Fisher Scientific), 0.5% MEM-NEAA (Sigma Aldrich), 1 % Penicillin/Streptomycin, 0.025% Insulin solution (Sigma Aldrich) and 2g/l Bicarbonate (Sigma Aldrich) and organoids were moved to orbital shakers in 6 or 10 cm dishes. Organoids were fed twice a week. 40 d after organoid generation, organoids were either cultured in HN+A or were transferred gradually to Low-Nutrient medium (LN) consisting of BrainPhys Neuronal Medium (Stem Cell Technologies), 2% B27+A (50X, Thermo Fisher Scientific), 1% N2 supplement (Thermo Fisher Scientific), 200 nM Ascorbic Acid (Sigma Aldrich), 0.2% CD Lipid Concentrate (Thermo Fisher Scientific), 1% Penicillin/Streptomycin, 1% Matrigel (Corning)^4^. In addition, the glucose concentration of LN was adjusted to 10 mM and the following supplements were added freshly before use: 20 ng/ml BDNF (Stem Cell Technologies), 20 ng/ml GDNF (Stem Cell Technologies) and 1 mM db-cAMP (Santa Cruz). Gradual transfer from HN+A to LN was performed as follows with feedings performed every third day, thus the total transition time was 12 days: 1. HN+A 75%:25% LN, 2. HN+A 50%:50% LN, 3. HN+A 25%:75% LN, 4. HN+A 0%:100% LN.

### Treatment of organoids with Everolimus and Afatinib

Organoids were treated with 20 nM of Everolimus (Abcam) as described^5^. A stock solution of 10 μM was prepared in DMSO (Sigma Aldrich) and diluted 1:500 in medium. Afatinib (Selleckchem) was used at a final concentration of 1 μM as published^6^. A stock solution of 500 μM was prepared in DMSO (Sigma Aldrich) and diluted 1:500 in medium. DMSO was used as a control and diluted 1:500 in medium. DMSO, Everolimus or Afatinib were freshy added to High Nutrient Medium and used to feed organoids from day 110 on. Organoids were fed with medium containing freshly added inhibitors every three days. After 30 days of treatment (d140) organoids were fixed and analyzed.

### FACS sorting

EGFR+ cells were isolated using Flow cytometry. Briefly, organoids were harvested, washed with DPBS−/− (Thermo Fisher Scientific) and added to a gentleMACS dissociator C tube (Miltenyi). 2 ml of Trypsin (Sigma Aldrich)/Accutase (Sigma Aldrich) (1:1) was added containing 10 U/ml DNaseI (Thermo Fisher Scientific). Organoids were dissociated using the 37C_NTDK_1 program on a gentleMACS™ Octo Dissociator with Heaters (Miltenyi). After dissociation, cells were briefly spun down to collect the sample, passed through a 70 μm cell strainer (Falcon) and 8 ml of PBS with 10% FBS was added. Cells were spun at 400×g for 4 min and supernatant was removed. Cells were resuspended in 2 ml PBS with 10% FBS, counted and 5 μL/10^6^ cells of Human EGFR (Research Grade Cetuximab Biosimilar) Alexa Fluor^®^ 488-conjugated Antibody (R&D Systems) was added. Cells were incubated for 30 min on ice and washed 3 times with 1 ml of PBS with 10% FBS. Cells were then passed through a 35 μm cell strainer and sorted on a SH800 Cell Sorter (Sony).

### Immunohistochemistry on paraffin embedded human samples

Use of human brain samples for histological analysis was approved by the local ethics committee of the Medical University of Vienna. Human brain material was processed for routine histopathology. Tissue was formalin fixed and embedded in paraffin and 3μm thin FFPE tissue sections were prepared. Immunohistochemistry for EGFR, TSC2, MAP2 and GFAP was performed using the Bond III automatic stainer (Leica) using the conditions described (Extended Data Table 1). Immunohistochemistry for Vimentin, GFAP and Secretagogin was performed using the EnVision™ FLEX+ kit (Dako) as detection system and diaminobenzidine (DAB) as chromogen with the conditions described (Extended Data Table 1). The histological staining procedure occurred with the help of coverplates (Thermo Fisher Scientific, Glass Coverplates). All sections were counterstained with haematoxylin.

### Immunohistochemistry on frozen human brain samples

Tissue was collected with consent in strict observance of the legal and institutional ethical regulations of the University of California San Francisco Committee on Human Research. Protocols for use of prenatal human tissues were approved by the Human Gamete, Embryo and Stem Cell Research Committee (institutional review board) at the University of California, San Francisco and postnatal autopsy and surgical tissues were obtained via the UCSF Pediatric Neuropathology Research Laboratory and Brain Tumor Research Center. Brains were fixed in 4% paraformaldehyde for 2 days, and then cryoprotected in a 30% sucrose solution. Blocks were cut into 30-micron sections on a cryostat and mounted on glass slides for immunohistochemistry. Frozen slides were allowed to equilibrate to room temperature for 3 hours. 10 minutes antigen retrieval was conducted at 95°C in 10 mM Na Citrate buffer, pH=6.0. Following antigen retrieval, slides were washed with TNT (0.05% TX100 in PBS) for 10 minutes, placed in 1% H2O2 in PBS for 90 minutes, and then blocked with TNB solution (0.1 M Tris-HCl, pH 7.5, 0.15 M NaCl, 0.5% blocking reagent from PerkinElmer) for 1 hour. Slides were incubated in primary antibodies overnight at 4°C (Extended Data Table 1) and in biotinylated secondary antibodies (Jackson Immunoresearch Laboratories) for 2.5 hours at room temperature. All antibodies were diluted in TNB solution from PerkinElmer. Sections were then incubated for 30 min in streptavidin-horseradish peroxidase that was diluted (1:200) with TNB. Tyramide signal amplification (Perkin-Elmer) was used for some antigens. Sections were incubated in tyramide-conjugated fluorophores at the following dilutions: Fluorescein, 1:50 (4.5 min); Opal 570, 1:100 (10 min); Cy5, 1:50 (4.5 min). Sections were imaged on a Leica TCS SP8 microscope.

### Immunohistochemistry on frozen organoid samples

Immunohistochemistry was performed as described^7,8^. Briefly, organoids were fixed in 4% paraformaldehyde for 20 - 40 min at room temperature. Organoids were washed with PBS three times for 10 min each at room temperature and then allowed to sink in 30% sucrose at 4 °C. Organoids were embedded in 7.5% gelatin (Sigma Aldrich) in 10% sucrose (Sigma Aldrich) solution and sectioned at 20 μm on a Cryostat (Leica NX70). For immunohistochemistry, sections were blocked and permeabilized in 0.25% TritonX-100 with 4% normal donkey serum (Millipore) for 1 hour at room temperature. Sections were then incubated at least 5 hours to overnight at room temperature with primary antibodies in 0.1% Triton X-100 with 4% normal donkey serum (Millipore) in PBS. Sections were washed 3 times in PBS. Secondary antibodies were incubated for 1hr and afterwards sections were washed 3 times in PBS. DAPI was added to secondary antibody to mark nuclei. Secondary antibodies labeled with Alexafluor 488, 568, or 647 (Invitrogen) were used for detection. Slides were mounted using Fluorescent mounting medium (Dako).

All antibodies used have been validated for the performed application as evidenced by validation profile on Antibodypedia, or manufacturer’s website.

### Laser microdissection

Organoids were fixed in 4% paraformaldehyde for 20 min at room temperature. Organoids were washed with PBS three times for 10 min each at room temperature and then allowed to sink in 30% sucrose (Sigma Aldrich) at 4 °C. Organoids were then embedded in 7.5% gelatin (Sigma Aldrich) in 10% sucrose solution and sectioned at 20 μm on a Cryostat (Leica NX70). Consecutive slides were mounted on glass slides and immunohistochemistry was performed to identify tumors or mounted on Frame Slides for microdissection (Leica; 11505190). Tumor regions were identified by EGFR and pS6 staining on consecutive slides and corresponding regions were dissected using the LMD6500 laser microdissection system (Leica). DNA was isolated from dissected regions using the DirectPCR^®^ Lysis Reagent Tail (Peqlab) containing 0.375 mg/ml Proteinase K (Thermo Fisher Scientific) according to the manufacturer’s instructions.

### Microscopy

Images were acquired on LSM880 and LSM800 confocal laser scanning microscopes (Zeiss). Large overview images were acquired using the Pannoramic Slide Scanner 250 Flash II or III system (3DHistech).

### Quantification of immunofluorescence

Quantifications of areas of dysmorphic neurons and giant cells were performed using ImageJ Software. For dysmorphic neurons wild type references were generated by analyzing regions of 6 organoids from 2 batches for each isogenic control line. For *TSC2*^+/−^ organoids 5 regions of about 500μm x 500μm with dysmorphic neurons were analyzed. 20 to 40 dysmorphic neurons were analyzed per organoid (Ext. Data Fig. 3F). Only after analysis of the size in Map2 staining, the expression of COUP-TFII and SCGN was analyzed.

For giant cells, size of pS6 expressing cells in one region of roughly 650μm x 650μm per organoid was analyzed. Whole organoids were acquired with the Pannoramic Slide Scanner 250 Flash II or III system (3DHistech) and then regions exported using the CaseViewer Software (3DHistech). Size analysis was performed in ImageJ. For isogenic Ctrl representative regions were chosen randomly. For *TSC2*^+/−^ organoids regions with giant cells were chosen. Evident by the analysis also normal sized pS6 cells were analyzed in *TSC2*^+/−^ organoids.

For the analysis of TSC2 expression in giant cells one region of giant cells per organoid was selected in whole organoid scans with the Pannoramic Slide Scanner 250 Flash II or III system (3DHistech). Expression of TSC2 was evaluated in visually enlarged appearing giant cells, as the size differences can be clearly distinguished by eye.

For quantifications of tumor areas, whole organoid sections were acquired with the Pannoramic Slide Scanner 250 Flash II or III system (3DHistech) and exported to ImageJ. Areas with double positive staining for Ki67 and pS6 were measured using the circle tool.

Quantifications of tumor areas for treatment experiments were performed using the CaseViewer Software (3DHistech). Briefly, whole organoid sections were acquired with the Pannoramic Slide Scanner 250 Flash II or III system (3DHistech). Areas with double positive staining for pS6 and EGFR were measured using the circle tool.

### Single cell RNA sequencing

Organoids were harvested, washed with DPBS−/− and incubated in Trypsin (Sigma Aldrich)/Accutase (Sigma Aldrich) (1:1) containing 10 U/ml DNaseI (Thermo Fisher Scientific) at 37°C on a thermoblock with agitation (800 rpm) for 40-60 min. A half volume of ice-cold DPBS−/− (Thermo Fisher Scientific) with 0.04% BSA (Sigma-Aldrich) was added to the dissociated cells. The solution was passed twice through a 35 μm cell strainer and centrifuged for 5 min at 400g. Cells were washed with once with DPBS−/− (Thermo Fisher Scientific) with 0.04% BSA (Sigma Aldrich) and resuspended in 150 μl ice-cold PBS containing 0.04% BSA (Sigma Aldrich). Cells were counted and 16000 cells were loaded per channel (to give estimated recovery of 10,000 cells per channel) onto a Chromium Single Cell 3′ B Chip (10x Genomics, PN-1000073) and processed through the Chromium controller to generate single-cell GEMs (Gel Beads in Emulsion). scRNA-seq libraries were prepared with the Chromium Single Cell 3′ Library & Gel Bead Kit v.3 (10x Genomics, PN-1000075). Libraries were pooled and sequenced paired end (R1:26, R2:98 cycles) on a NextSeq 550 (Illumina) at 200 million reads per library.

### Single-cell RNA-seq data analysis

In our analysis we largely followed a recently published approach^9^. Firstly, we aligned reads to GRCh38 human reference genome with Cell Ranger 3.0 (10× Genomics) using default parameters to produce the cell-by-gene, Unique Molecular Identifier (UMI) count matrix. UMI counts were analyzed using the Seurat R package v.3. Cells were filtered for a min. 1000 & max. 6000 genes, maximal mitochondrial content of 15%, and maximal ribosomal content of 30%. Resulting high quality cells were normalized (“LogNormalize”) for scaled for each cell to a total expression of 10K UMI. Variable genes were identified by Seurat’s FindVariableFeatures implementation (“FastLogVMR”). We found that regression for UMI count, mitochondrial or ribosomal content (via ScaleData) does not improve the separation of known cell types, therefore we did not regress out any known variables.

### Data integration and Batch correction

We integrated the 4 sequencing libraries (batches 1 & 2 for wt & TSC2) using Canonical Correlation Analysis (CCA), based on integration anchors calculated on the first 30 components. Briefly, correlation vectors across datasets are aligned by “dynamic time warping” algorithm, resulting in all cells embedded in a shared low-dimensional CCA space, where batch variation is minimized. Cell-to-cell distances in aligned CCA space are the basis for later visualization (UMAP).

### Clustering

For Principal Components Analysis (PCA), the data was transformed (‘ScaleData’) so that the mean expression of each gene across cells is 0, and that the variance is 1, ensuring equal weight to each gene, regardless of absolute expression level. Principal components were calculated on the variable genes. We then calculated the k-nearest neighbors of each cell from Euclidean distances in PCA-space (using top 30 principal components). Next, we constructed the shared nearest neighbor (SNN) graph, where each edge (between two cells) is weighted by the fraction of overlap between the two cells’ k-nearest neighbors (also known as Jaccard similarity), using default parameters. Louvain clustering on this graph (resolution=0.3) identified 16 clusters. Clusters were visualized on UMAPs. UMAP is a nonlinear dimensionality reduction technique compressing cell-to-cell distances calculated in CCA space to two 2 (plotting) dimensions.

### Cluster identity

We calculated differentially expressed genes in each cluster (compared to the rest of the cells) using Seurat’s Wilcoxon test implementation and analyzed genes with Bonferroni-adjusted P value < 0.05, along the expression of classical marker genes of know cell types. Based on this, we manually labeled each cell type (Fig.2A).

### Integration of Organoid and fetal single-cell data

To confirm the identity of our cell types, we integrated our organoid data with recently published single-cell data from fetal brains of various developmental ages^10^. The fetal samples have been sequenced on the same platform (10X Chromium) and were obtained from https://cells.ucsc.edu. The fetal dataset was integrated with our organoid data the same way as described above, with the following differences: The fetal datasets from different individuals showed large variation in quality, with the following fractions passing the above defined quality thresholds: week 6: 97% = 5970 cells; week 10: 93% = 7193 cells; week 14: 18% = 14435 cells; week 18: 93% = 78157 cells; week 22: 32% = 83619 cells. Such strongly skewed distribution of high-quality cells towards late embryos (week 18 & 22) would skew the integrated data analysis, where the variance identified would mostly be representative of late development. To achieve a more equal representation of different developmental stages, we (1) relaxed the minimum UMI filtering criteria to 800 and the maximal ribosomal content to 40%, plus (2) we down sampled datasets (post filtering) to similar numbers of cells (max 10000 cells per fetal sample, and max 5000 per organoid). The integrated analysis otherwise followed the steps described above. Transcriptome-wide for figure 2D, the average gene expression of CLIP-cells from organoids is compared to fetal cells’ average gene expression by Pearson correlation coefficients.

### Analysis of whole genome sequencing data

Samples were processed using the nfcore/sarek pipeline v2.5.1^11^, more specifically reads were aligned to GRCh38 using bwa mem v0.7.17, alignments were post-processed using GATK v4.1.2.0 according to GATK best practices, ascatNgs was used to perform genome-wide allele-specific copy number analysis. A circos visualization of the overall genomic variant profile was generated with hmftools (purple v2.34, cobalt v1.7, amber v3.0) using somatic variations identified by Strelka v2.9.10.

### Mass spectrometry

#### Sample preparation

Cell pellets for the FACS sorting were processed via iST kit 96× (PreOmics GmbH) according original protocol from manufacturer. Briefly, pellet was mixed with 50 μl of Lysis buffer and incubated 10 min in 95 °C. After cooling to room temperature lysate was mixed with 50 μl of digest solution and incubated overnight in 37 °C. In the next step solution was transferred into cartridge with 100 μl of Stop solution. After the washing by solution Wash1 and Wash2 (each 200 μl) peptides were eluted from the cartridge by Elute solution in two steps (each with 100 μl). Peptide solution was completely dried via Speed Vac.

Dried peptides were solubilized in 50 μl of 0.1 % TFA and sonicate in ultrasonication bath Sonorex RK52 (Bandelin). Peptide solution was stored in −80°C prior the nanoLC-MS/MS analysis.

#### Stable isotope labelled (SIL) peptides synthesis

To determine retention time of targeted peptides of TSC1 and TSC2 stable isotope labelled peptides were synthetized. N-terminal amino acid was changed for the [^13^C_6_,^15^N_2_]-lysine respectively for the [^13^C_6_,^15^N_4_]-arginine. Peptides were synthetized using solid-phase Fmoc chemistry, purified using preparative reversed-phase chromatography, lyophilized, and subsequently characterized by MALDI-TOF-MS (using an ABI 4800 MALDI-TOF/TOF, SCIEX Peptides were pooled and spiked in real samples before the injection into nanoLC-MS/MS system.

#### NanoLC-MS/MS analysis and data processing

The nano HPLC system used was an UltiMate 3000 RSLC nano system coupled to a Q Exactive HF-X mass spectrometer, equipped with the with an EASY-Spray™source (Thermo Fisher Scientific) and Jailbreak 1.0 (Phoenix S&T). Peptides were loaded onto a trap column (Thermo Fisher Scientific, PepMap C18, 5 mm × 300 μm ID, 5 μm particles, 100 Å pore size) at a flow rate of 25 μL/min using 0.1% TFA as mobile phase. After 10 min, the trap column was switched in line with the analytical column (Thermo Fisher Scientific, PepMap C18, 500 mm × 75 μm ID, 2 μm, 100 Å). Peptides were eluted using a flow rate of 230 nl/min, and a binary 2h gradient, respectively 165 min.

The gradient starts with the mobile phases: 98% A (water/formic acid, 99.9/0.1, v/v) and 2% B (water/acetonitrile/formic acid, 19.92/80/0.08, v/v/v), increases to 35% B over the next 130 min, followed by a gradient in 5 min to 90% B, stays there for 5 min and decreases in 2 min back to the gradient 98% A and 2% B for equilibration at 30°C.

The Q Exactive HF-X mass spectrometer was operated by a mixed MS method which consisted of one full scan (*m/z* range 380-1,500; 15,000 resolution; target value 1e6) followed by the PRM of targeted peptides from an inclusion list (isolation window 0.7 *m/z*; normalized collision energy (NCE) 30; 30,000 resolution, AGC target 2e5). The maximum injection time variably changed based on the number of targets in the inclusion list to use up the total cycle time of 600 ms. The scheduling window were set to 4 min for each precursor.

List of peptides including basic mass spectrometry information used for PRM analysis of TSC1, TSC2 and 5 normalization proteins are displayed in the table in Ext. Data Figure 13.

Data processing and manual evaluation of results were performed in Skyline-daily (64-bit, v19.0.9.190.^12^). For the data processing peptides were used which had at least 3 specific peptide fragments. TSC1 and TSC2 proteins were quantified based on integrated ion intensities over retention time of peptides from inclusion list. To account for different amounts between samples, these values were normalized based on a set of five abundant/house-keeping proteins (SRP14, LMNB1, HIST1H1B, GAPDH and MDH2). Based on the abundance of these proteins, normalization factors for each normalization protein and sample were computed. The median of these factors per sample was used to normalize TSC1 and TSC2 abundance.

## References

1 Lui, J. H., Hansen, D. V. & Kriegstein, A. R. Development and evolution of the human neocortex. Cell 146, 18–36, doi:10.1016/j.cell.2011.06.030 (2011).

2 Pollen, A. A. et al. Establishing Cerebral Organoids as Models of Human-Specific Brain Evolution. Cell 176, 743–756 e717, doi:10.1016/j.cell.2019.01.017 (2019).

3 Nowakowski, T. J. et al. Spatiotemporal gene expression trajectories reveal developmental hierarchies of the human cortex. Science 358, 1318–1323, doi:10.1126/science.aap8809 (2017).

4 Hansen, D. V., Lui, J. H., Parker, P. R. & Kriegstein, A. R. Neurogenic radial glia in the outer subventricular zone of human neocortex. Nature 464, 554–561, doi:10.1038/nature08845 (2010).

5 Fietz, S. A. et al. OSVZ progenitors of human and ferret neocortex are epithelial-like and expand by integrin signaling. Nat Neurosci 13, 690–699, doi:10.1038/nn.2553 (2010).

6 Paredes, M. F. et al. Extensive migration of young neurons into the infant human frontal lobe. Science 354, doi:10.1126/science.aaf7073 (2016).

7 Henske, E. P., Jozwiak, S., Kingswood, J. C., Sampson, J. R. & Thiele, E. A. Tuberous sclerosis complex. Nat Rev Dis Primers 2, 16035, doi:10.1038/nrdp.2016.35 (2016).

8 Thiele, E. A. Managing and understanding epilepsy in tuberous sclerosis complex. Epilepsia 51 Suppl 1, 90–91, doi:10.1111/j.1528-1167.2009.02458.x (2010).

9 Ruppe, V. et al. Developmental brain abnormalities in tuberous sclerosis complex: a comparative tissue analysis of cortical tubers and perituberal cortex. Epilepsia 55, 539–550, doi:10.1111/epi.12545 (2014).

10 Curatolo, P., Bombardieri, R. & Jozwiak, S. Tuberous sclerosis. The Lancet 372, 657–668, doi:10.1016/s0140-6736(08)61279-9 (2008).

11 Crino, P. B. Evolving neurobiology of tuberous sclerosis complex. Acta Neuropathol 125, 317–332, doi:10.1007/s00401-013-1085-x (2013).

12 Knudson JR., A. G. Mutation and Cancer: Statistical Study of Retinoblastoma. Proc. Nat. Acad. Sci. USA Vol. 68, pp. 820–823,, doi:10.1073/pnas.68.4.820.

13 Feliciano, D. M., Su, T., Lopez, J., Platel, J. C. & Bordey, A. Single-cell Tsc1 knockout during corticogenesis generates tuber-like lesions and reduces seizure threshold in mice. J Clin Invest 121, 1596–1607, doi:10.1172/JCI44909 (2011).

14 Feliciano, D. M., Quon, J. L., Su, T., Taylor, M. M. & Bordey, A. Postnatal neurogenesis generates heterotopias, olfactory micronodules and cortical infiltration following single-cell Tsc1 deletion. Hum Mol Genet 21, 799–810, doi:10.1093/hmg/ddr511 (2012).

15 Way, S. W. et al. Loss of Tsc2 in radial glia models the brain pathology of tuberous sclerosis complex in the mouse. Hum Mol Genet 18, 1252–1265, doi:10.1093/hmg/ddp025 (2009).

16 Carson, R. P., Van Nielen, D. L., Winzenburger, P. A. & Ess, K. C. Neuronal and glia abnormalities in Tsc1-deficient forebrain and partial rescue by rapamycin. Neurobiol Dis 45, 369–380, doi:10.1016/j.nbd.2011.08.024 (2012).

17 Goto, J. et al. Regulable neural progenitor-specific Tsc1 loss yields giant cells with organellar dysfunction in a model of tuberous sclerosis complex. Proc Natl Acad Sci U S A 108, E1070–1079, doi:10.1073/pnas.1106454108 (2011).

18 Blair, J. D., Hockemeyer, D. & Bateup, H. S. Genetically engineered human cortical spheroid models of tuberous sclerosis. Nat Med 24, 1568–1578, doi:10.1038/s41591-018-0139-y (2018).

19 Qin, W. et al. Analysis of TSC cortical tubers by deep sequencing of TSC1, TSC2 and KRAS demonstrates that small second-hit mutations in these genes are rare events. Brain Pathol 20, 1096–1105, doi:10.1111/j.1750-3639.2010.00416.x (2010).

20 Henske, E. P. et al. Allelic Loss Is Frequent in Tuberous Sclerosis Kidney Lesions but Rare in Brain Lesions. Am J Hum Genet, 400–406 (1996).

21 Martin, K. R. et al. The genomic landscape of tuberous sclerosis complex. Nat Commun 8, 15816, doi:10.1038/ncomms15816 (2017).

22 Chan, J. A. et al. Pathogenesis of Tuberous Sclerosis Subependymal Giant Cell Astrocytomas: Biallelic Inactivation of TSC1 or TSC2 Leads to mTOR Activation. Journal of Neuropathology and Experimental Neurology (2004).

23 Lancaster, M. A. et al. Cerebral organoids model human brain development and microcephaly. Nature 501, 373–379, doi:10.1038/nature12517 (2013).

24 Bardy, C. et al. Neuronal medium that supports basic synaptic functions and activity of human neurons in vitro. Proc Natl Acad Sci U S A 112, E3312, doi:10.1073/pnas.1509741112 (2015).

25 Park, S. H. et al. Tuberous sclerosis in a 20-week gestation fetus: immunohistochemical study. Acta Neuropathol 94, 180–186 (1997).

26 Mizuguchi, M. & Takashima, S. Neuropathology of tuberous sclerosis. Brain & Development, 508–515 (2001).

27 Buccoliero, A. M. et al. Subependymal giant cell astrocytoma: a lesion with activated mTOR pathway and constant expression of glutamine synthetase. Clin Neuropathol 35, 295–301, doi:10.5414/NP300936 (2016).

28 Buccoliero, A. M. et al. Subependymal giant cell astrocytoma (SEGA): Is it an astrocytoma? Morphological, immunohistochemical and ultrastructural study. Neuropathology 29, 25–30, doi:10.1111/j.1440-1789.2008.00934.x (2009).

29 Velasco, S. et al. Individual brain organoids reproducibly form cell diversity of the human cerebral cortex. Nature 570, 523–527, doi:10.1038/s41586-019-1289-x (2019).

30 Bhaduri, A. et al. Cell stress in cortical organoids impairs molecular subtype specification. Nature, doi:10.1038/s41586-020-1962-0 (2020).

31 Baser, A. et al. Onset of differentiation is post-transcriptionally controlled in adult neural stem cells. Nature 566, 100–104, doi:10.1038/s41586-019-0888-x (2019).

32 LiCausi, F. & Hartman, N. W. Role of mTOR Complexes in Neurogenesis. Int J Mol Sci 19, doi:10.3390/ijms19051544 (2018).

33 Bulut-Karslioglu, A. et al. Inhibition of mTOR induces a paused pluripotent state. Nature 540, 119–123, doi:10.1038/nature20578 (2016).

34 Dulken, B. W., Leeman, D. S., Boutet, S. C., Hebestreit, K. & Brunet, A. Single-Cell Transcriptomic Analysis Defines Heterogeneity and Transcriptional Dynamics in the Adult Neural Stem Cell Lineage. Cell Rep 18, 777–790, doi:10.1016/j.celrep.2016.12.060 (2017).

35 Yuzwa, S. A. et al. Developmental Emergence of Adult Neural Stem Cells as Revealed by Single-Cell Transcriptional Profiling. Cell Rep 21, 3970–3986, doi:10.1016/j.celrep.2017.12.017 (2017).

36 Basak, O. et al. Troy+ brain stem cells cycle through quiescence and regulate their number by sensing niche occupancy. Proc Natl Acad Sci U S A 115, E610–E619, doi:10.1073/pnas.1715911114 (2018).

37 Shin, J. et al. Single-Cell RNA-Seq with Waterfall Reveals Molecular Cascades underlying Adult Neurogenesis. Cell Stem Cell 17, 360–372, doi:10.1016/j.stem.2015.07.013 (2015).

38 Parker, W. E. et al. Enhanced epidermal growth factor, hepatocyte growth factor, and vascular endothelial growth factor expression in tuberous sclerosis complex. Am J Pathol 178, 296–305, doi:10.1016/j.ajpath.2010.11.031 (2011).

39 Lim, J. S. et al. Somatic Mutations in TSC1 and TSC2 Cause Focal Cortical Dysplasia. Am J Hum Genet 100, 454–472, doi:10.1016/j.ajhg.2017.01.030 (2017).

40 Vinters, H. V. et al. Tuberous sclerosis-related gene expression in normal and dysplastic brain. Epilepsy Res 32, 12–23 (1998).

41 Johnson, M. W., Emelin, J. K., Park, S. H. & Vinters, H. V. Co-Localization of TSC1 and TSC2 Gene Products in Tubers of Patients with Tuberous Sclerosis. Brain Pathol, 45–54 (1999).

42 Franz, D. N. et al. Rapamycin causes regression of astrocytomas in tuberous sclerosis complex. Ann Neurol 59, 490–498, doi:10.1002/ana.20784 (2006).

43 Franz, D. N. et al. Everolimus for subependymal giant cell astrocytoma in patients with tuberous sclerosis complex: 2-year open-label extension of the randomised EXIST-1 study. The Lancet Oncology 15, 1513–1520, doi:10.1016/s1470-2045(14)70489-9 (2014).

44 Bissler, J. J. et al. Sirolimus for angiomyolipoma in tuberous sclerosis complex or lymphangioleiomyomatosis. N Engl J Med 358, 140–151, doi:10.1056/NEJMoa063564 (2008).

45 Krueger, D. A. et al. Long-term treatment of epilepsy with everolimus in tuberous sclerosis. Neurology 87, 2408–2415, doi:10.1212/WNL.0000000000003400 (2016).

46 Krueger, D. A. et al. Everolimus for subependymal giant-cell astrocytomas in tuberous sclerosis. N Engl J Med 363, 1801–1811, doi:10.1056/NEJMoa1001671 (2010).

47 Martins, F. et al. A review of oral toxicity associated with mTOR inhibitor therapy in cancer patients. Oral Oncol 49, 293–298, doi:10.1016/j.oraloncology.2012.11.008 (2013).

48 Kerfoot, C. et al. Localization of Tuberous Sclerosis 2 mRNA and its Protein Product Tuberin in Normal Human Brain and in Cerebral Lesions of Patients with Tuberous Sclerosis. Brain Pathol, 367–377 (1996).

49 Raju, C. S. et al. Secretagogin is Expressed by Developing Neocortical GABAergic Neurons in Humans but not Mice and Increases Neurite Arbor Size and Complexity. Cereb Cortex 28, 1946–1958, doi:10.1093/cercor/bhx101 (2018).

50 Hansen, D. V. et al. Non-epithelial stem cells and cortical interneuron production in the human ganglionic eminences. Nat Neurosci 16, 1576–1587, doi:10.1038/nn.3541 (2013).

51 Hodge, R. D. et al. Conserved cell types with divergent features in human versus mouse cortex. Nature 573, 61–68, doi:10.1038/s41586-019-1506-7 (2019).

52 Curatolo, P., Moavero, R. & de Vries, P. J. Neurological and neuropsychiatric aspects of tuberous sclerosis complex. The Lancet Neurology 14, 733–745, doi:10.1016/s1474-4422(15)00069-1 (2015).

## References

1 Agu, C. A. et al. Successful Generation of Human Induced Pluripotent Stem Cell Lines from Blood Samples Held at Room Temperature for up to 48 hr. Stem Cell Reports 5, 660–671, doi:10.1016/j.stemcr.2015.08.012 (2015).

2 Jinek, M. et al. A programmable dual-RNA-guided DNA endonuclease in adaptive bacterial immunity. Science 337, 816–821, doi:10.1126/science.1225829 (2012).

3 Lancaster, M. A. & Knoblich, J. A. Generation of cerebral organoids from human pluripotent stem cells. Nat Protoc 9, 2329–2340, doi:10.1038/nprot.2014.158 (2014).

4 Bardy, C. et al. Neuronal medium that supports basic synaptic functions and activity of human neurons in vitro. Proc Natl Acad Sci U S A 112, E3312, doi:10.1073/pnas.1509741112 (2015).

5 Blair, J. D., Hockemeyer, D. & Bateup, H. S. Genetically engineered human cortical spheroid models of tuberous sclerosis. Nat Med 24, 1568–1578, doi:10.1038/s41591-018-0139-y (2018).

6 Bian, S. et al. Genetically engineered cerebral organoids model brain tumor formation. Nat Methods 15, 631–639, doi:10.1038/s41592-018-0070-7 (2018).

7 Lancaster, M. A. et al. Cerebral organoids model human brain development and microcephaly. Nature 501, 373–379, doi:10.1038/nature12517 (2013).

8 Lancaster, M. A. et al. Guided self-organization and cortical plate formation in human brain organoids. Nat Biotechnol 35, 659–666, doi:10.1038/nbt.3906 (2017).

9 Velasco, S. et al. Individual brain organoids reproducibly form cell diversity of the human cerebral cortex. Nature 570, 523–527, doi:10.1038/s41586-019-1289-x (2019).

10 Bhaduri, A. et al. Cell stress in cortical organoids impairs molecular subtype specification. Nature, doi:10.1038/s41586-020-1962-0 (2020).

11 Garcia, M. et al. Sarek: A portable workflow for whole-genome sequencing analysis of germline and somatic variants. F1000Research 9, doi:10.12688/f1000research.16665.1 (2020).

12 MacLean, B. et al. Skyline: an open source document editor for creating and analyzing targeted proteomics experiments. Bioinformatics 26, 966–968, doi:10.1093/bioinformatics/btq054 (2010).

