## Supplemental Figures for "Cerebral organoid model reveals excessive proliferation of human caudal late interneuron progenitors in Tuberous Sclerosis Complex"

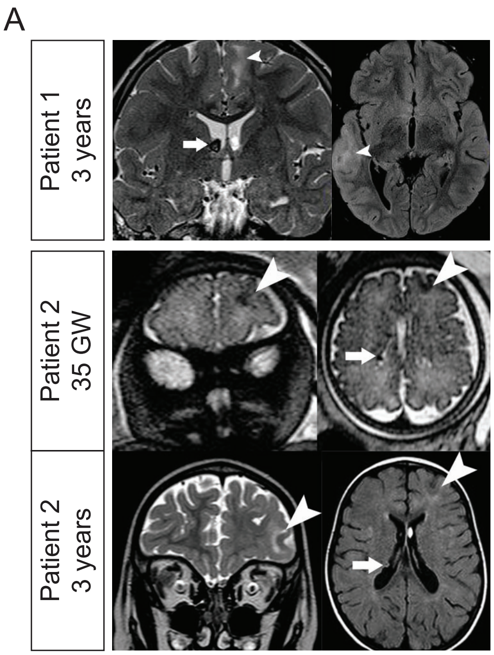


B

| | DOB | Tubers | SEGA | Seizure onset | seizure onset zone | current seizure frequency | epilepsy surgery | neuropsych. Impairment |
|---|---|---|---|---|---|---|---|---|
| Patient 1 | August 2005 | yes | yes | 7 months | left frontal, fronto-parietal | 4 per day (partially psych.) | no | high |
| Patient 2 | July 2007 | yes | yes | 6 months | left frontal | auras 0.5/day | left frontal (2010) | mild |

C

| | Previous AEDs | Current AEDs |
|---|---|---|
| Patient 1 | LAM, LEV, OX, Top, VAL, VGA, GABA, FORT, Zon, Fycompa | Everolimus (2016), Zonesamid, perampanel, midazolam (during seizure) |
| Patient 2 | VGB, VPA, LEV | Everolimus |

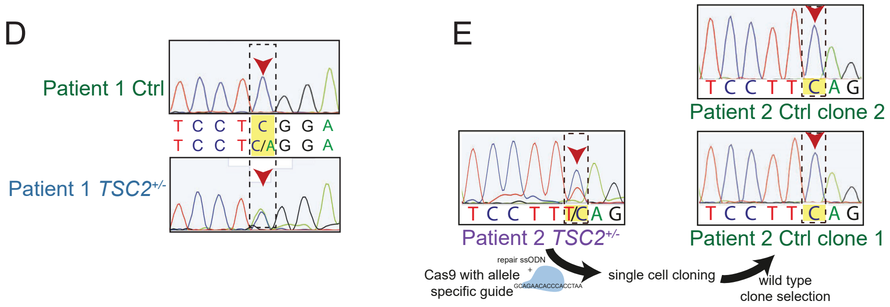


**Extended Data Fig. 1**

**A.** MRI images of patients included in this study. Top panel: Patient 1 MRI at 3 years of age shows cortical tuber (arrowhead) and subependymal tumor (arrow). Middle panel: Fetal MRI image of Patient 2 at 35GW shows presence of tubers (arrowhead) and tumors (arrow), already prenatally. Bottom panel: At 3 years of age the tubers and tumors are still detected.

**B.** Background information on Patients and clinical presentations. Both patients show tubers and tumors as well as drug-resistant epilepsy and neuropsychiatric impairment.

**C.** Antiepileptic drugs (AEDs) used previously and currently in patients reprogrammed in this study.

**D.** Sequencing information of Patient 1. Reprogramming of Patient 1 yielded heterozygous mutant, as well as homozygous wild type clones, as this patient is mosaic for the disease-causing mutation.

**E.** Sequencing information of Patient 2. Reprogramming resulted in heterozygous clones, of which one was repaired with purified Cas9 protein and allele specific guide adding a repair single-stranded-oligodeoxynucleotide (ssODN) as HDR repair template. Two clones were selected and followed up during this study.

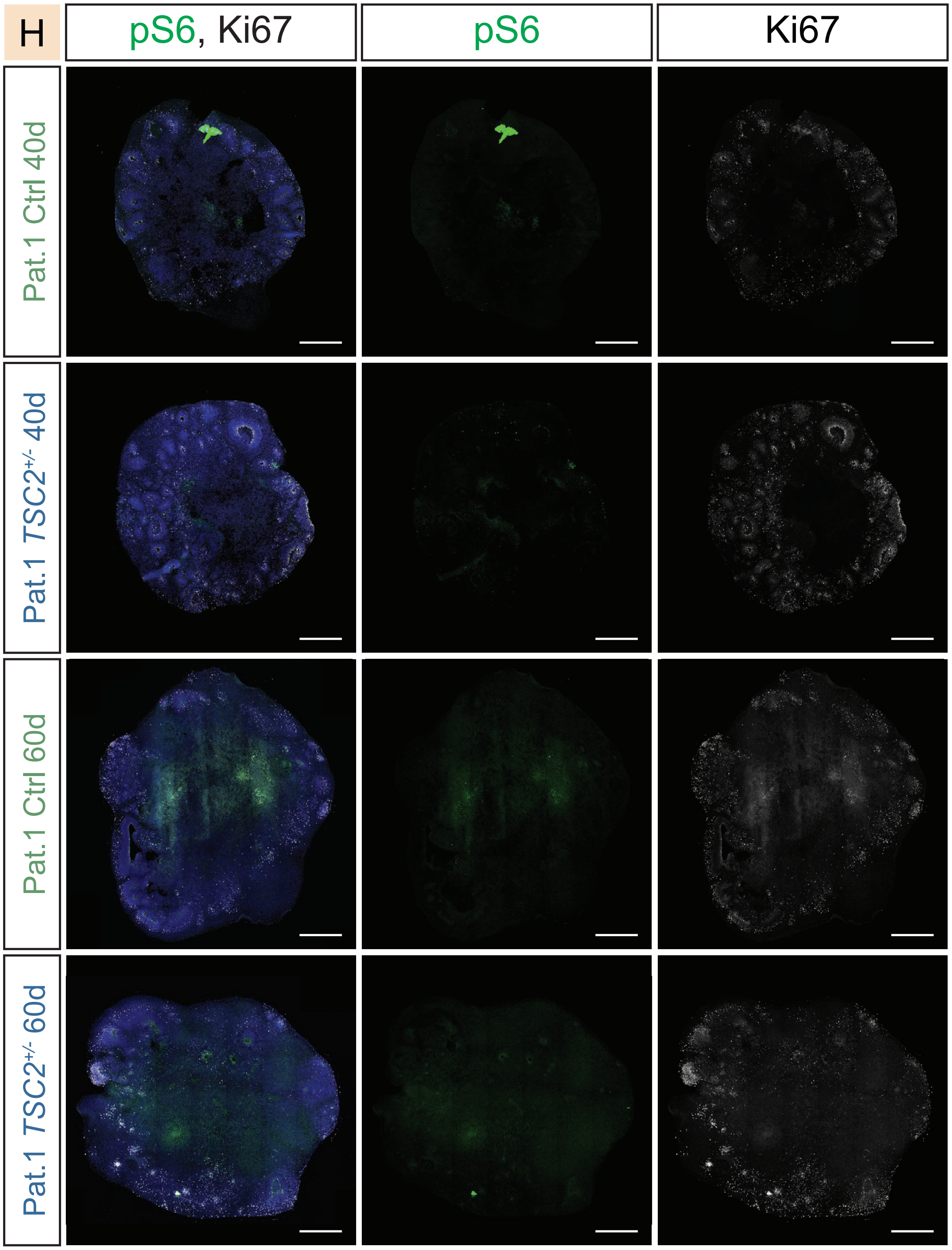


**Ext. Data Fig. 2 – Early Ctrl or *TSC2*$^{+/-}$-derived organoids show no differences**
pS6 and Ki67 staining shows no difference between Ctrl or *TSC2*$^{+/-}$-derived organoids at 40 and 60 days after EB formation.
(Scale bar: 500µm)

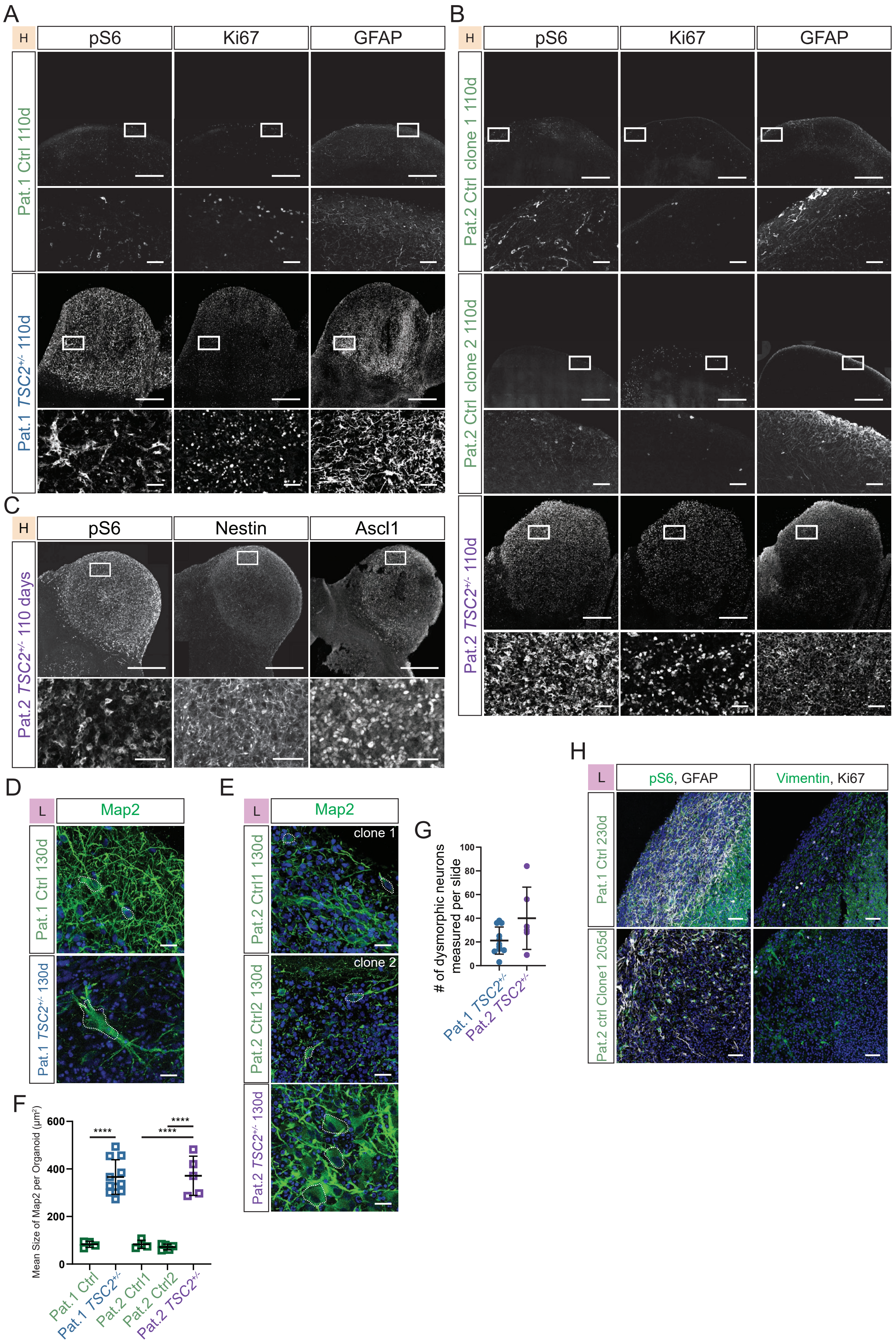
A
H
pS6
Ki67
GFAP
Pat.1 Ctrl 110d
Pat.1 TSC2+/- 110d
B
H
pS6
Ki67
GFAP
Pat.2 Ctrl clone 1 110d
Pat.2 Ctrl clone 2 110d
Pat.2 TSC2+/- 110d
C
H
pS6
Nestin
Ascl1
Pat.2 TSC2+/- 110 days
D
L
Map2
Pat.1 Ctrl 130d
Pat.1 TSC2+/- 130d
E
L
Map2
clone 1
clone 2
Pat.2 Ctrl1 130d
Pat.2 Ctrl2 130d
Pat.2 TSC2+/- 130d
F
Mean Size of Map2 per Organoid (µm²)
600
400
200
0
****
****
****
Pat.1 Ctrl
Pat.1 TSC2+/-
Pat.2 Ctrl1
Pat.2 Ctrl2
Pat.2 TSC2+/-
G
### of dysmorphic neurons measured per slide
100
80
60
40
20
0
Pat.1 TSC2+/-
Pat.2 TSC2+/-
H
L
pS6, GFAP
Vimentin, Ki67
Pat.1 Ctrl 230d
Pat.2 ctrl Clone1 205d

**Ext. Data Fig. 3 – *TSC2*$^{+/-}$-derived organoids show proliferative tumors marked by pS6 and Ki67 at 110 days**
**A.** and **B.** Immunostainings for pS6, Ki67 and GFAP on Pat.1 ctrl, Pat.1 *TSC2*$^{+/-}$, two repaired clones of Pat.2 and Pat.2 *TSC2*$^{+/-}$-derived organoids. Isogenic controls show individual cells with pS6 staining, but no nodular proliferative tumors. *TSC2*$^{+/-}$-derived organoids show nodular aggregations of pS6 positive and Ki67-expressing proliferative nodules. Morphology as well as expression of GFAP resembles what was previously described in TSC patient SEN/SEGAs.
**C.** SEN-like tumors in organoids express the NSC markers Nestin and Ascl1.
**D.** and **E.** Immunostaining for Map2 identifies enlarged dysmorphic neurons *TSC2*$^{+/-}$-derived organoids in low-nutrient medium. In the isogenic controls, neurons have normal size and barely cytoplasm are stained.
**F.** Size of Map2 in *TSC2*$^{+/-}$ dysmorphic and Ctrl normal neurons. The mean of the measured size of all neurons per organoid in µm2 is shown (Pat.1 Ctrl N=2, n=4 mean= 83, SD=11.2; Pat.1 *TSC2*$^{+/-}$ N=6, n=11, 241 dysmorphic neurons, mean=366.1, SD=69.4, Pat.1 Ctrl vs. *TSC2*$^{+/-}$ p<0.0001; Pat.2 Ctrl1 N=2, n=4, 433 neurons, mean=83, SD=15; Pat.2 Ctrl2 N=2, n=5, 597 neurons, mean=71.7, SD=10.7; Pat.2 *TSC2*$^{+/-}$ N=2, n=6, 240 dysmorphic neurons, mean=371.3, SD=73.9, Pat.2 Ctrl vs. *TSC2*$^{+/-}$ and Pat.2 Ctrl vs. *TSC2*$^{+/-}$ both p<0.0001; test: Ordinary One-Way ANOVA)
**G.** Number of dysmorphic neurons measured per slide of *TSC2*$^{+/-}$-derived organoids. As no dysmorphic neurons appear in Ctrl organoids only *TSC2*$^{+/-}$ samples are shown. Always around 20-40 neurons were measured per slide (Pat.1 *TSC2*$^{+/-}$ N=6, n=11, 241 dysmorphic neurons, mean=21.18, SD=10.9, Pat.2 *TSC2*$^{+/-}$ N=2, n=6, 240 dysmorphic neurons, mean=40, SD=24).
**H.** Immunostainings for pS6, GFAP and Vimentin, Ki67 on isogenic control organoids at 230 days in L-medium. GFAP staining identifies individual cell bodies and processes. Some cell bodies also stain for pS6, but are much smaller than giant cells in *TSC2*$^{+/-}$ organoid (Fig. 1F). Vimentin staining identifies individual normal sized cells and proliferative marker Ki67 shows a low proliferation.
(Scale bars: overview images A. and B.: 500µm; insets A., B. and C.: 50µm; C. and D.: 20µm)

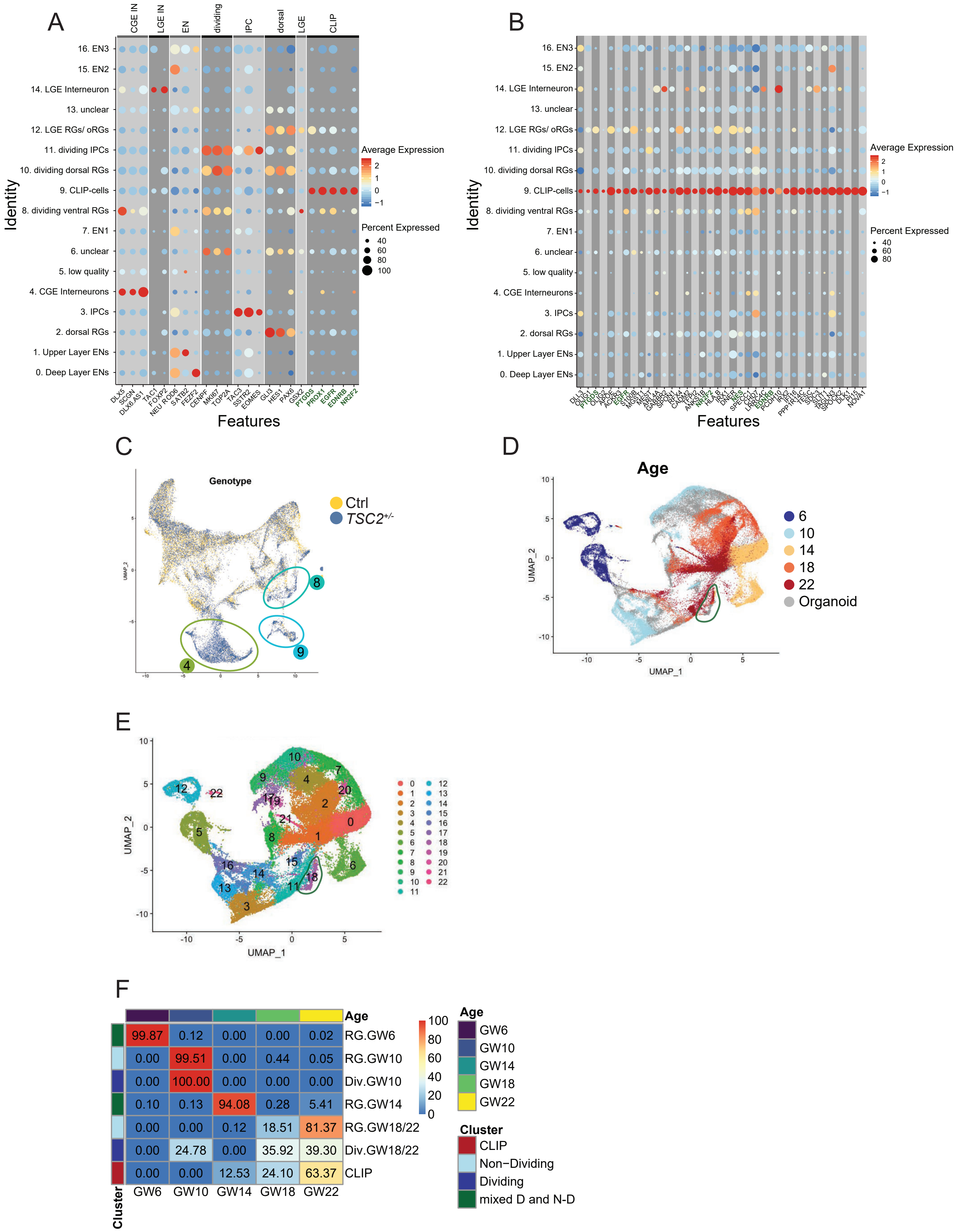

A
CGE IN
LGE IN
EN
dividing
IPC
dorsal
LGE
CLIP
16. EN3
15. EN2
14. LGE Interneuron
13. unclear
12. LGE RGs/ oRGs
11. dividing IPCs
10. dividing dorsal RGs
9. CLIP-cells
8. dividing ventral RGs
7. EN1
6. unclear
5. low quality
4. CGE Interneurons
3. IPCs
2. dorsal RGs
1. Upper Layer ENs
0. Deep Layer ENs
Identity
DLX5 SCGN DLX6-AS1 TAC1 FOXP2 NEUROD6 SATB2 FEZF2 CENPF MKI67 TOP2A TAC3 SSTR2 EOMES GLI3 HES1 PAX6 GSX2 PTGDS PROX1 EGFR EDNRB NR2F2
Features
Average Expression
2
1
0
−1
Percent Expressed
40
60
80
100
B
Identity
Features
Average Expression
Percent Expressed
C
Genotype
Ctrl
TSC2+/-
UMAP_2
8
9
4
D
Age
6
10
14
18
22
Organoid
UMAP_1
UMAP_2
E
UMAP_1
UMAP_2
F
Age
Cluster
RG.GW6
RG.GW10
Div.GW10
RG.GW14
RG.GW18/22
Div.GW18/22
CLIP
GW6 GW10 GW14 GW18 GW22
99.87 0.12 0.00 0.00 0.02
0.00 99.51 0.00 0.44 0.05
0.00 100.00 0.00 0.00 0.00
0.10 0.13 94.08 0.28 5.41
0.00 0.00 0.12 18.51 81.37
0.00 24.78 0.00 35.92 39.30
0.00 0.00 12.53 24.10 63.37
100
80
60
40
20
0
Age
GW6
GW10
GW14
GW18
GW22
Cluster
CLIP
Non−Dividing
Dividing
mixed D and N-D

**Ext. Data Fig. 4 – scRNA-seq on Ctrl and *TSC2*$^{+/-}$-derived organoids**

**A.** DotPlot of marker genes for the different clusters. Size of the circle represents the percentage of cells within each cluster that express the respective gene. Expression in minimum 20% of the cells is required to draw a circle. Color-coded from blue to red, the average expression within the cluster is shown with a maximal scaled average expression of 2.5. Markers shown for CLIP-cells are written in bold green and have been used for validation.

**B.** DotPlot showing CLIP-cell specific genes. Differential gene expression analysis was performed against all other clusters, as well as pairwise against cluster 12 as CLIP-cells share a lot of genes with LGE RGs and oRGs. Same cut-offs as for the DotPlot in panel A were used. Markers used for validation of CLIP-cells are written in bold green.

**C.** UMAP projection showing the two genotypes color-coded. Note an increase in ventral dividing, as well as CGE-derived interneurons and CLIP-cells in *TSC2*$^{+/-}$ organoids.

**D.** UMAP projection of our organoid datasets merged with the primary cells from a recently published dataset (Bhaduri and Andrews 2020). Color coded the age of the primary cells is shown. Note that CLIP-cells (green circle) are mainly co-clusering with radial glia from later gestational ages.

**E.** UMAP projection of different clusters integrating organoid and primary dataset. CLIP-cells cluster in cluster 18 (green circle).

**F.** Heatmap showing the relative cluster contributions of samples fromdifferent gestational ages to each cluster. Relative abundance of cells from each age was computed on cell numbers downsampled to the smallest dataset. Clusters were defined based on gene expression identifying radial glia and dividing radial glia clusters for GW10 and later samples (GW18/22). For GW6 and GW14 both non-dividing and dividing are in one cluster. Most cells contributing to the CLIP-cell cluster originate from GW22.

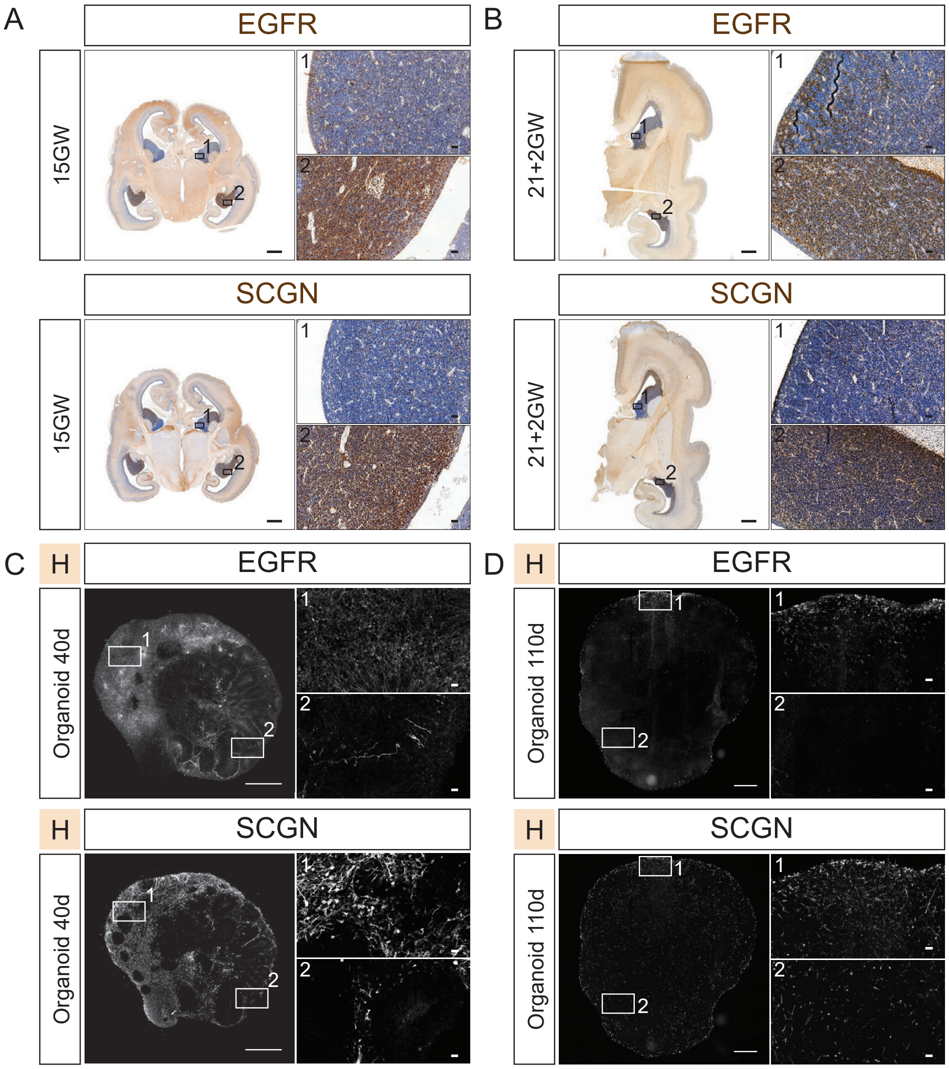


**Ext. Data Fig. 5 – EGFR is expressed in the developing CGE**

**A.** EGFR is expressed in CGE regions marked by SCGN around 15 gestational weeks. Insets show EGFR in CGE region (1) and negative MGE (2).

**B.** Similarly, EGFR expression is found in CGE regions in during later periods of gestation. Insets show EGFR in CGE region (1) and negative MGE (2).

**C.** Top panel: Early organoids express EGFR in CGE-patterned regions marked by SCGN (Inset 1), while expression in other regions is low (Inset 2).

**D.** EGFR expression is still found in Ctrl organoids at later age in regions that also show SCGN expression, indicating EGFR continues to be expressed in CGE-patterned regions in a later developmental stage in organoids. Insets show CGE-patterned region co-expressing EGFR and SCGN region not expressing both markers.

(Scale Bars: overview images A. and B.: 2mm; overview images C. and D.: 500µm; Insets A. to D.: 50µm)

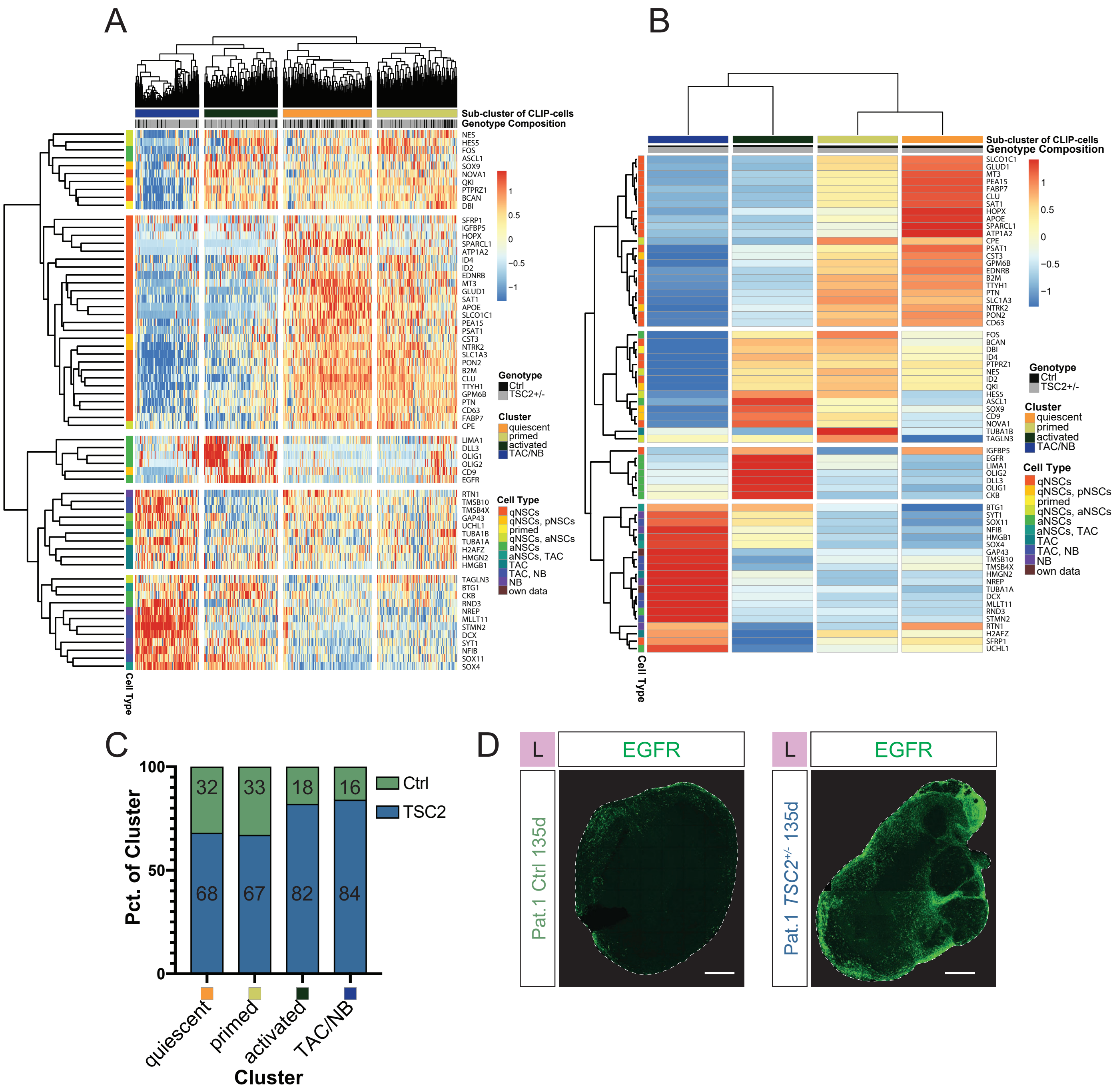


**Ext. Data Fig. 6 – Heatmap Clustering of CLIP-cells**

**A.** Heatmap of hierachical clustering of CLIP-cells showing scaled expression per cell. A minimal RNA expression of 2 as well as expression in minimal 30 cells was set as thresholds for genes. Genes were clustered with correlation distance and cells with manhattan distance. Note that in contrast to average expression for each group of cells, as shown in Fig. 2F and Ext. Data Fig 5B, the genes are cut into 5 groups with two TAC/NB modules. As in Fig. 2F color-coded on the left the annotation of each gene is shown ranging from quiescent over primed and activated to TAC/NB associated genes with genes shared in multiple states. Note that in contrast to Fig. 2F the order of the gene modules is: Primed, quiescent, activated, and two TAC/NB modules. On top the different clusters of cells are shown: TAC/NB, activated, quiescent and primed from left to right. Below color coded in gray and black the genotype of each cell is shown.

**B.** Heatmap as shown in Fig. 2F with annotated genes.

**C.** Genotype composition of each heatmap cluster showing activated clusters are mostly made up of *TSC2*$^{+/-}$ cells.

**D.** Immunostaining for EGFR, which is specifically expressed in activated CLIP-cells,  on whole organoids cultured in L-medium shows that EGFR is increased in *TSC2*$^{+/-}$ organoids.

Scale Bar D: 500µm

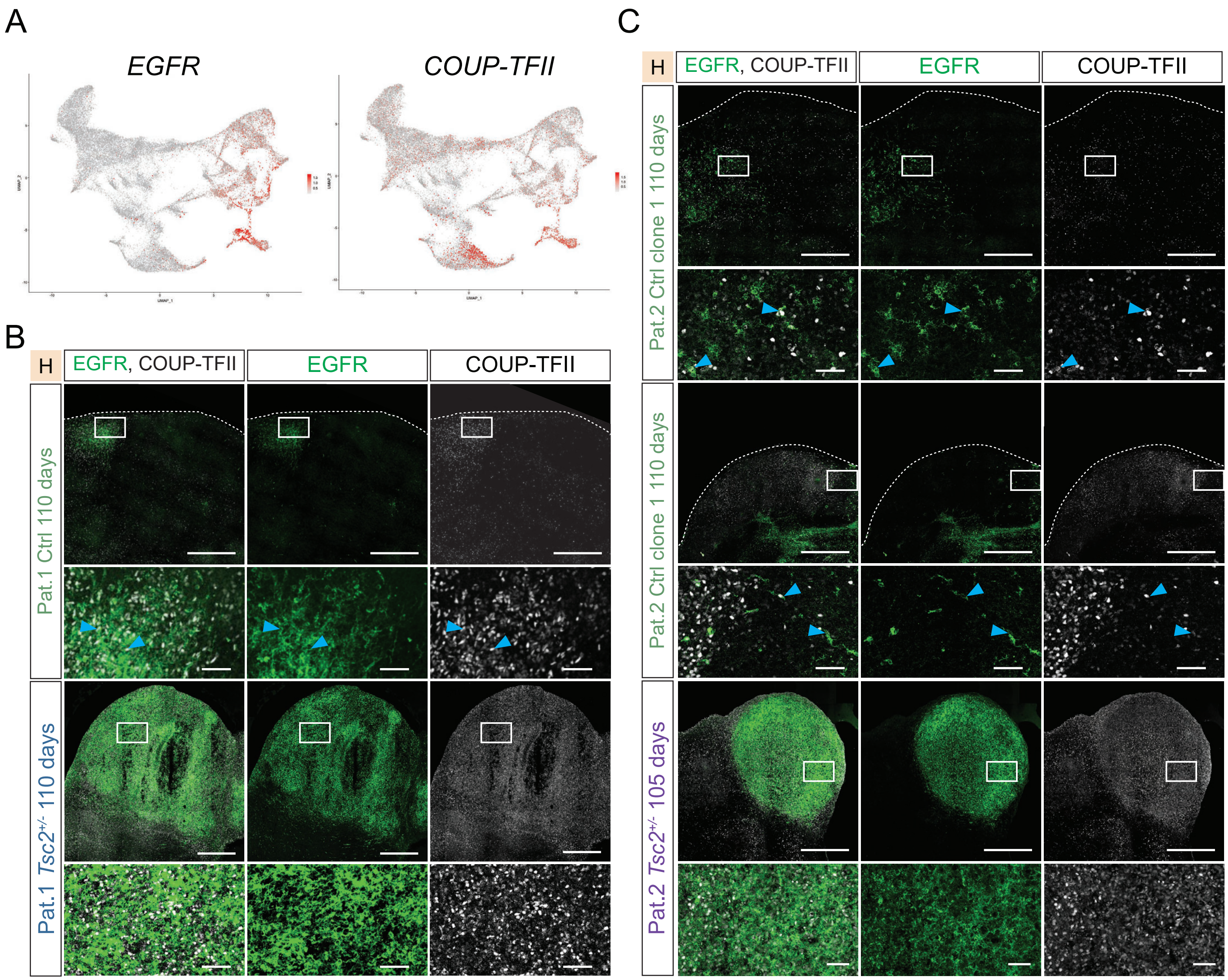


**Ext. Data Fig. 7 - EGFR and COUP-TFII specifically mark CLIP-cells in the scRNA-seq dataset and are expressed in tumors in *TSC2*$^{+/-}$-derived organoids**

**A.** Expression of *EGFR* and *COUP-TFII* on UMAP as in Fig. 2A. *EGFR* is specifically expressed in CLIP-cells, as well as in dividing cells of the ventral lineage. There is a low expression in dorsal progenitors. *COUP-TFII* expression in all cells showing specific expression in CLIP-Cells and interneurons.

**B.** and **C.** Validation of findings in scRNA-seq. Top: EGFR and COUP-TFII are co-expressed in individual cells in Ctrl organoids in Patient 1 and both repair clones of Patient 2 cultured in high-nutrient medium. Bottom: Previously identified tumors in *TSC2*$^{+/-}$-derived organoids are composed of EGFR and COUP-TFII positive cells in both patients. Arrowhead in insets of Ctrl organoids show co-expression of COUP-TFII and EGFR.

(Scale bars: overview images B. and C.: 500µm; Insets B. and C.: 50µm)

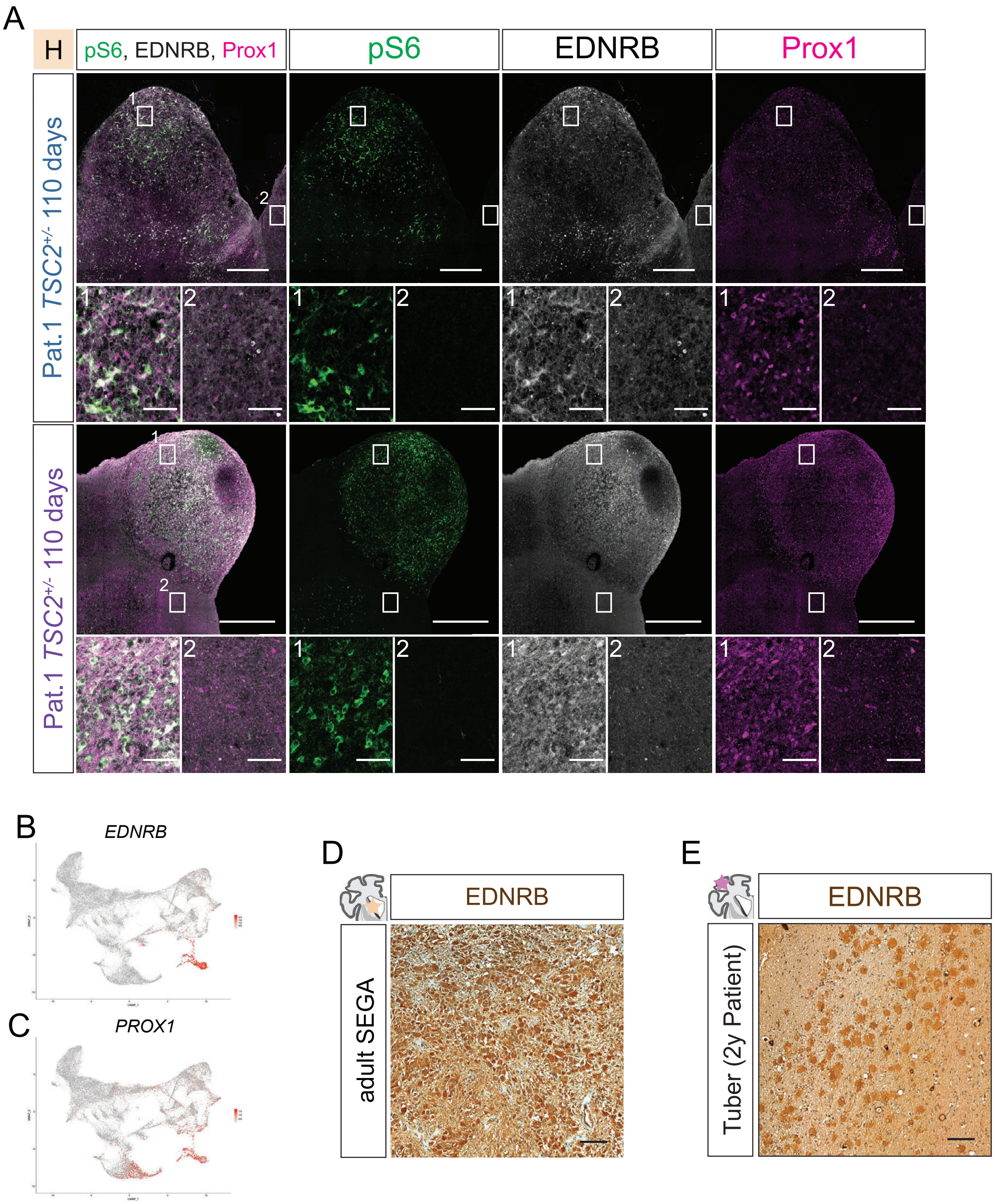


**Ext. Data Fig. 8 - EDNRB and Prox1 specifically mark CLIP-cells in the scRNA seq and are expressed in tumor cells in *TSC2*$^{+/-}$-derived organoids**

**A.** pS6 is co-expressed with EDNRB and Prox1 in tumors in *TSC2*$^{+/-}$-derived organoids. Insets 1 mark regions within the tumor, showing co-expression of all three markers. Insets 2 show regions outside of the tumor and are negative for all three markers. Note that pS6-high cells within tumor also express high levels of EDNRB.

**B.** *EDNRB* is a unique marker of CLIP-cells within scRNA-seq of all organoids (UMAP as in Fig. 2A).

**C.** *PROX1* is expressed in CLIP-cells and interneurons.

**D.** EDNRB expression in tumor cells is confirmed in adult resected SEGA .

**E.** Giant cells in a resected tuber from a 2 years-old patient express EDNRB.

(Scale bars: overview images A.: 500µm; Insets A.: 50µm, D. and E.: 100µm)

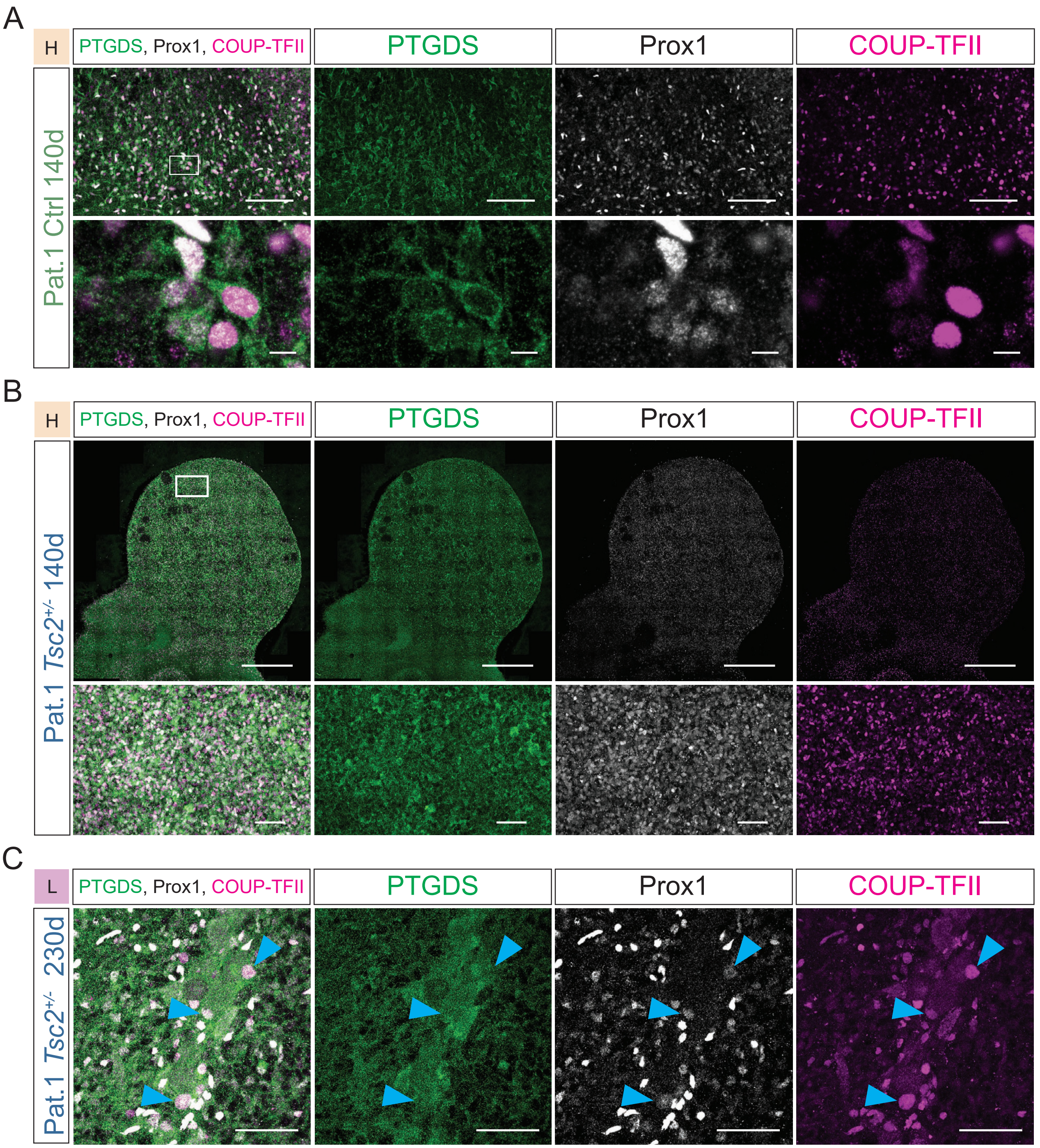


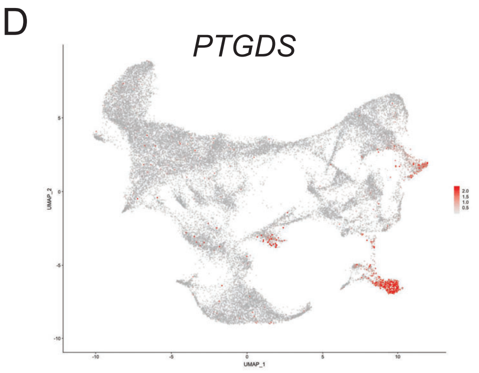


**Ext. Data Fig. 9 - Prostaglandin D Synthase together with Prox1 and COUP-TFII specifically labels CLIP-cells**

**A.** CLIP-cells in Ctrl organoids. Combination of PTGDS, Prox1 and COUP-TFII in 140d old organoids of Pat.1 Ctrl cultured in high-nutrient medium stains radial glia-like cells.

**B.** PTGDS, Prox1 and COUP-TFII are expressed in 140d-old organoids in tumors in high-nutrient medium.

**C.** PTGDS, Prox1 and Coup-TFII are expressed in 230d-old organoids in giant cells in low-nutrient medium.

**D.** *PTGDS* is expressed specifically in CLIP-cell cluster (UMAP as in Fig. 2A). See Ext. Data Fig 7 and 8 for *PROX1* and *COUP-TFII* expression in the same cluster.

(Scale bars: overview image A.: 100µm; Inset A.: 10µm; Overview image B.: 500µm; Inset B. and C.: 50µm)

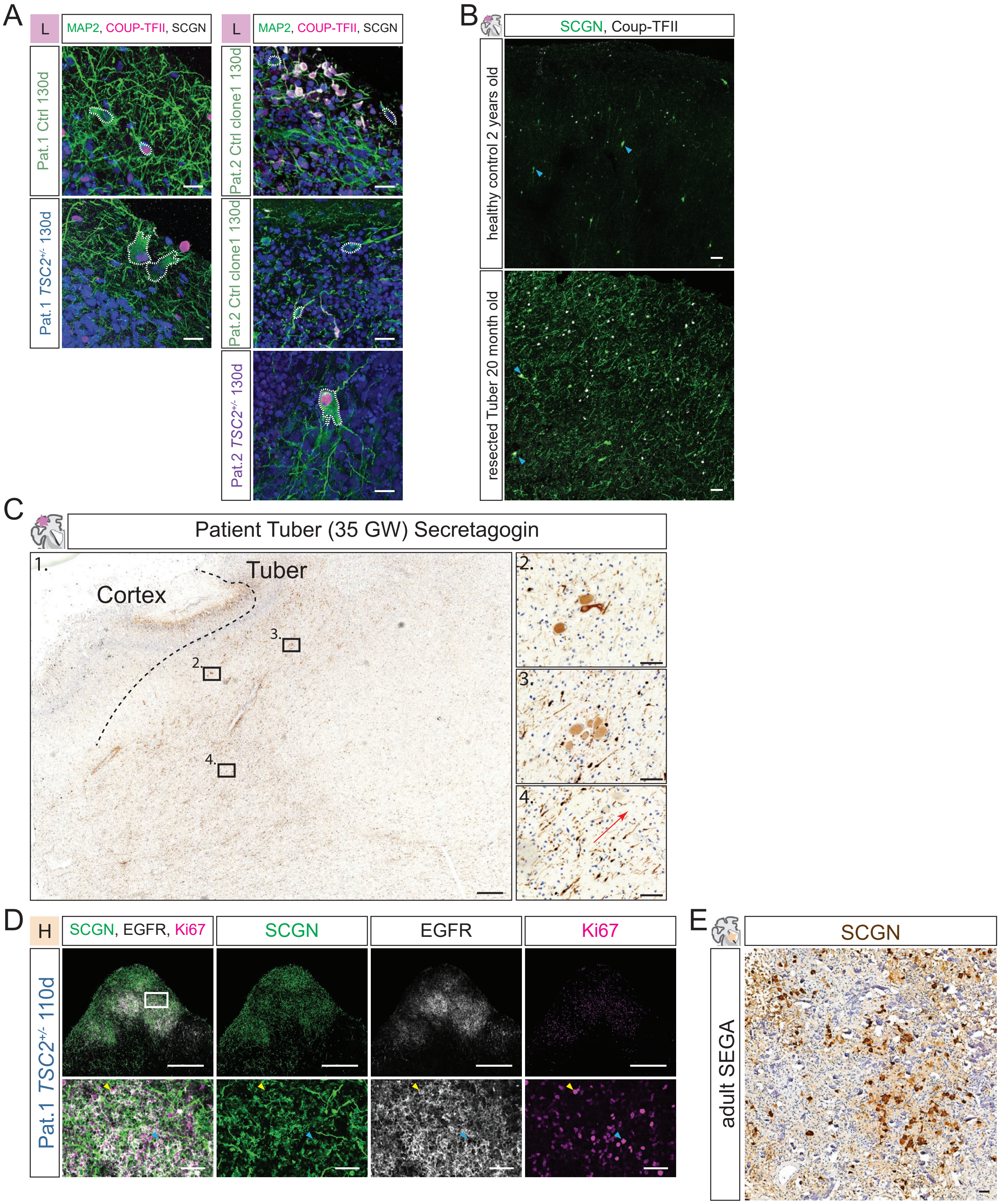

A
L
MAP2, COUP-TFII, SCGN
Pat.1 Ctrl 130d
Pat.1 TSC2+/- 130d
Pat.2 Ctrl clone1 130d
Pat.2 TSC2+/- 130d
B
SCGN, Coup-TFII
healthy control 2 years old
resected Tuber 20 month old
C
Patient Tuber (35 GW) Secretagogin
Cortex
Tuber
D
H
SCGN, EGFR, Ki67
SCGN
EGFR
Ki67
Pat.1 TSC2+/- 110d
E
SCGN
adult SEGA

**Ext. Data Fig. 10 - CGE-derived interneurons are found in TSC lesions**
**A.** MAP2, COUP-TFII and SCGN staining in 130 days-old organoids cultured in L-medium. Top: Isogenic Ctrls of Patient 1 (left) and 2 (right) show small individual neurons. Bottom: Dysmorphic neurons in *TSC2*$^{+/-}$-derived organoids of both patients express CGE-associated markers.
**B.** Overview of SCGN and COUP-TFII expression in the cortex of two years-old healthy control and in a resected tuber of a 20 months-old TSC patient. In the healthy control case individual neurons expressing SCGN with barely visible processes are stained. In contrast, in the resected tuber strong immunolabelling for both SCGN and COUP-TFII is detected in enlarged dysmorphic neurons (arrowheads). Additionally, thickened and dysmorphic processes throughout the whole tuber are strongly stained.
**C.** Secretagogin staining on formalin-fixed paraffin-embedded (FFPE) material of 35GW-old fetal TSC case. White matter is in left lower corner of overview image (1.). Grey matter is on the upper part of the overview, with the beginning of tuber from middle to right top (dashed line). Note enlarged dysmorphic neurons staining for SCGN in insets 2. and 3. at border and within tuber respectively. In 4., SCGN positive cells with migrating morphology within white matter can be identified. The majority of the leading processes indicating directionality points towards tuber (arrow).
**D.** SCGN, EGFR and Ki67 staining on tumor in 110 days-old organoid cultured in high-nutrient medium shows that SCGN positive cells can be generated in tumors. Note that there are SCGN, EGFR, Ki67 triple positive (blue arrowhead), as well as EGFR, Ki67 double positive cells (yellow arrowhead).
**E.** SCGN staining on FFPE material of resected adult SEGA. Enlarged SCGN positive cells can be identified. Differences in morphology compared to early tumors in organoids are likely due to disease progression in adult tumors.
(Scale bars: A.: 20µm; B., Insets C., Insets D. and E.: 50µm; overview C. and overview D.: 500µm)

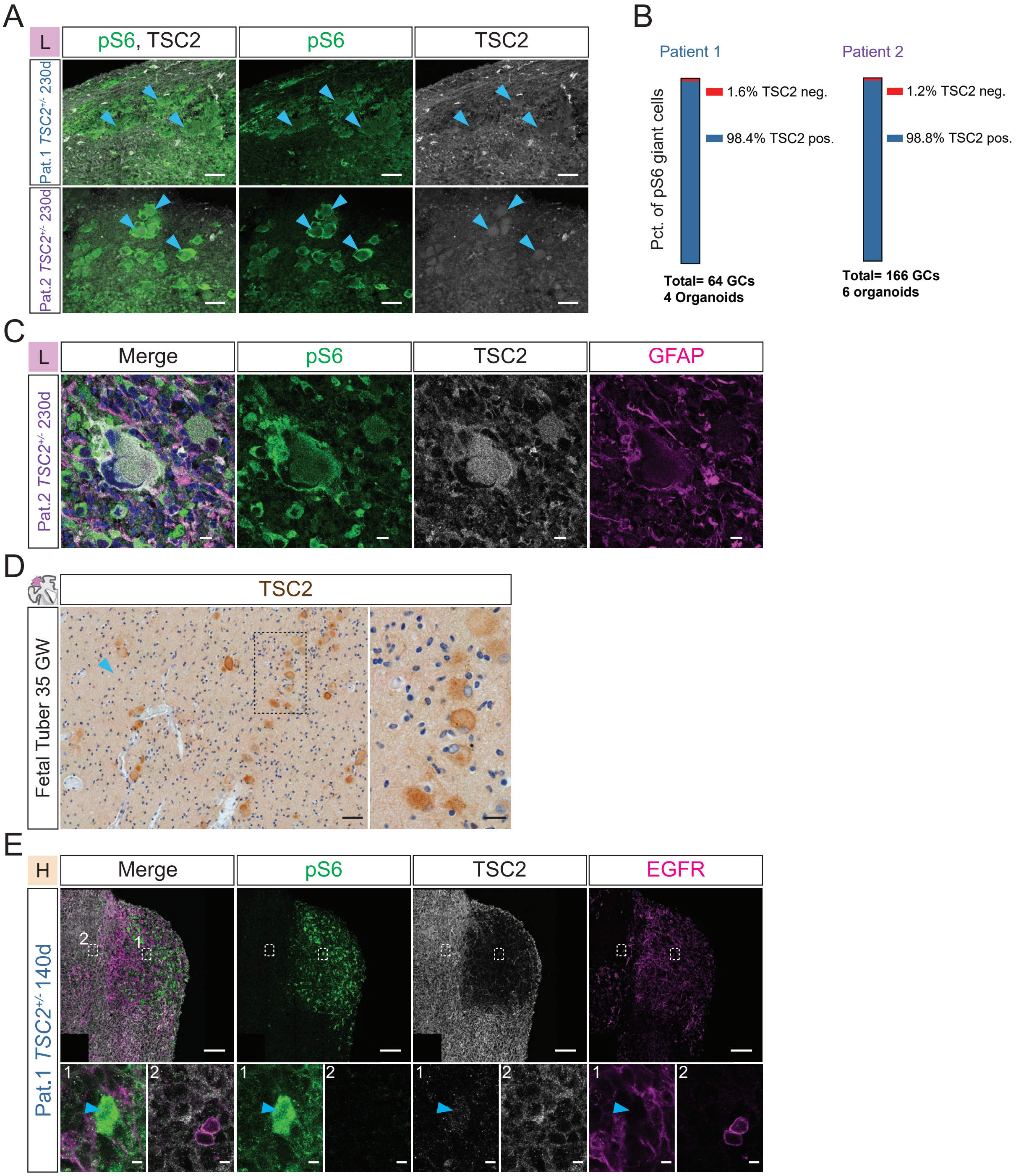


**Ext. Data Fig. 11 - TSC2 is expressed in giant cells but downregulated in tumors**
**A.** pS6 and TSC2 staining in tuber-like regions in 230 days-old organoids cultured in L-medium shows expression of TSC2 in pS6 positive Giant Cells.
**B.** Quantification of TSC2 expression in Giant cells shows almost all Giant cells in organoids show TSC2 expression (Pat.1 N=2, n=4, 64 giant cells; Pat.2 N=3, n=6, 166 giant cells).
**C.** High-Magnification of pS6 and GFAP expressing giant cell that co-expresses TSC2 in 230 days-old organoid.
**D.** TSC2 staining on 35GW-old fetal tuber confirms that TSC2 is expressed in tubers.
**E.** TSC2 staining on tumors in organoids shows focal reduction of TSC2. Note that pS6-positive cells within tumor weakly express TSC2, while the rest of the tumor is negative (blue arrow, inset 1). TSC2 is detected outside of tumor (inset 2).
(Scale bars: A. and overview D.: 50μm; C.: 10μm; D. inset: 20μm; E. overview: 100μm; E. inset: 5μm)

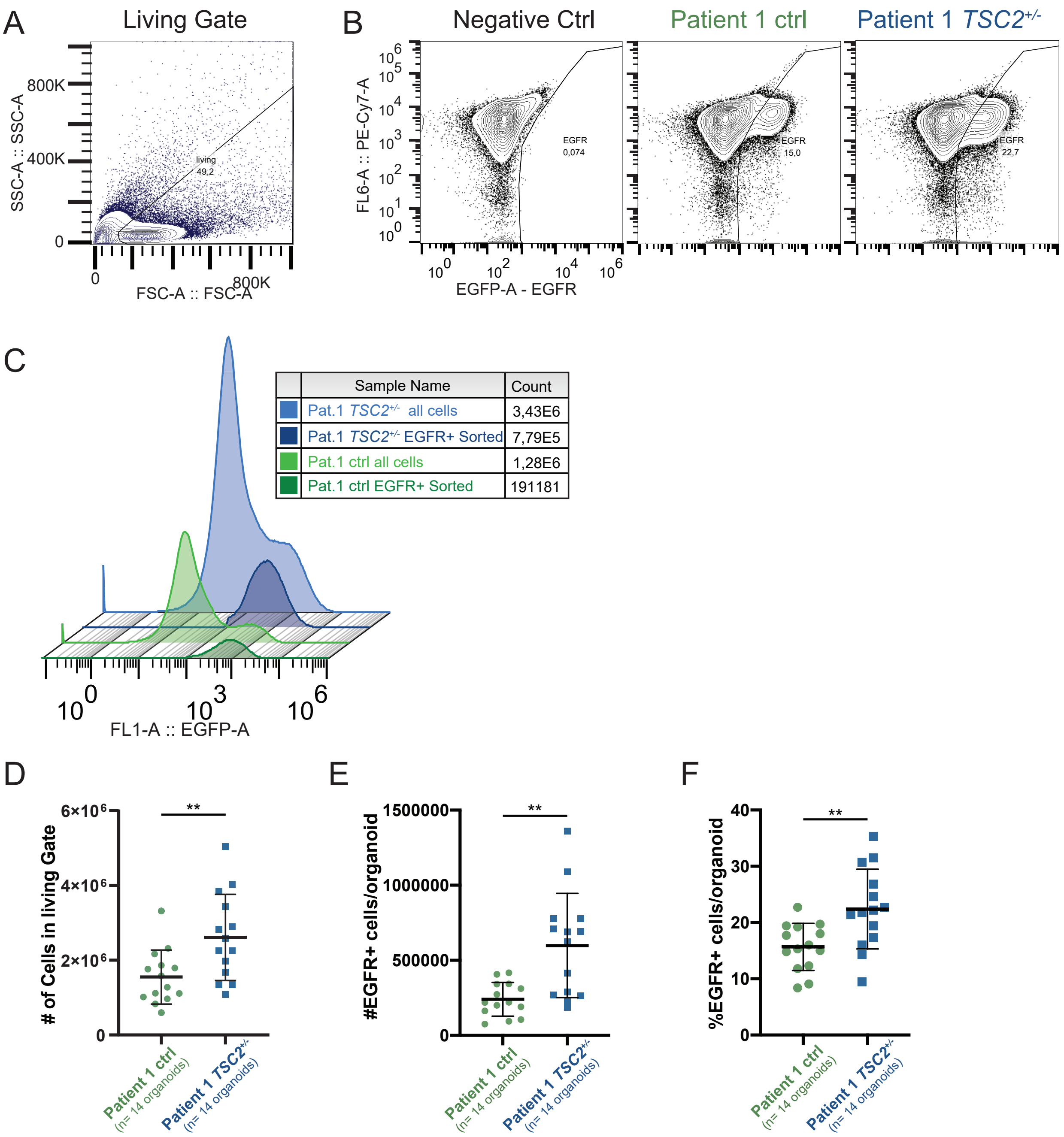


**Ext. Data Fig. 12 - FACS sorting of Pat.1 Ctrl and Pat.1 *TSC2*$^{+/-}$-derived tumor organoids**

**A.** Exemplary FACS plot of live cell gate on FSC-A and SSC-A.

**B.** Exemplary FACS plots with gating for EGFRpositive cells on unstained (left), Pat.1 Ctrl (middle) and Pat.1 *TSC2*$^{+/-}$-derived (right) single cell suspension.

**C.** Histogram of FACS plots shown in B with cell counts of all cells and EGFR+ sorted cells for Pat.1 Ctrl and Pat.1 *TSC2*$^{+/-}$-derived organoids. *TSC2*$^{+/-}$-derived organoids show more cells in total as well as relatively and absolutely more EGFR+ cells than Ctrl.

**D.** Total number of cells in living gate for all Ctrl- and *TSC2*$^{+/-}$-derived organoids shows mutant organoids have more cells in total (N=2, n=14 for both genotypes; mean Ctrl = 1.5E6 cells, SD = 0.7E6; mean TSC2+/- = 2.6E6 cells, SD = 1.1E6; p=0.0072, unpaired t-test).

**E.** Number of EGFR+ cells per genotype shows *TSC2*$^{+/-}$-derived organoids have more EGFR+ cells (N=2, n=14 for both genotypes; mean Ctrl = 2.4E5 cells, SD = 1.1E5; mean *TSC2*$^{+/-}$ = 5.98E5 cells, SD = 3.4E5; p=0.0011, unpaired t-test)

**F.** Percentage of EGFR+ cells per genotype. *TSC2*$^{+/-}$-derived organoids have higher percentage of EGFR+ cells (N=2, n=14 for both genotypes; mean Ctrl = 15.67%, SD = 4.02%; mean *TSC2*$^{+/-}$ = 22.39%, SD = 6.82%; p=0.0051, unpaired t-test).

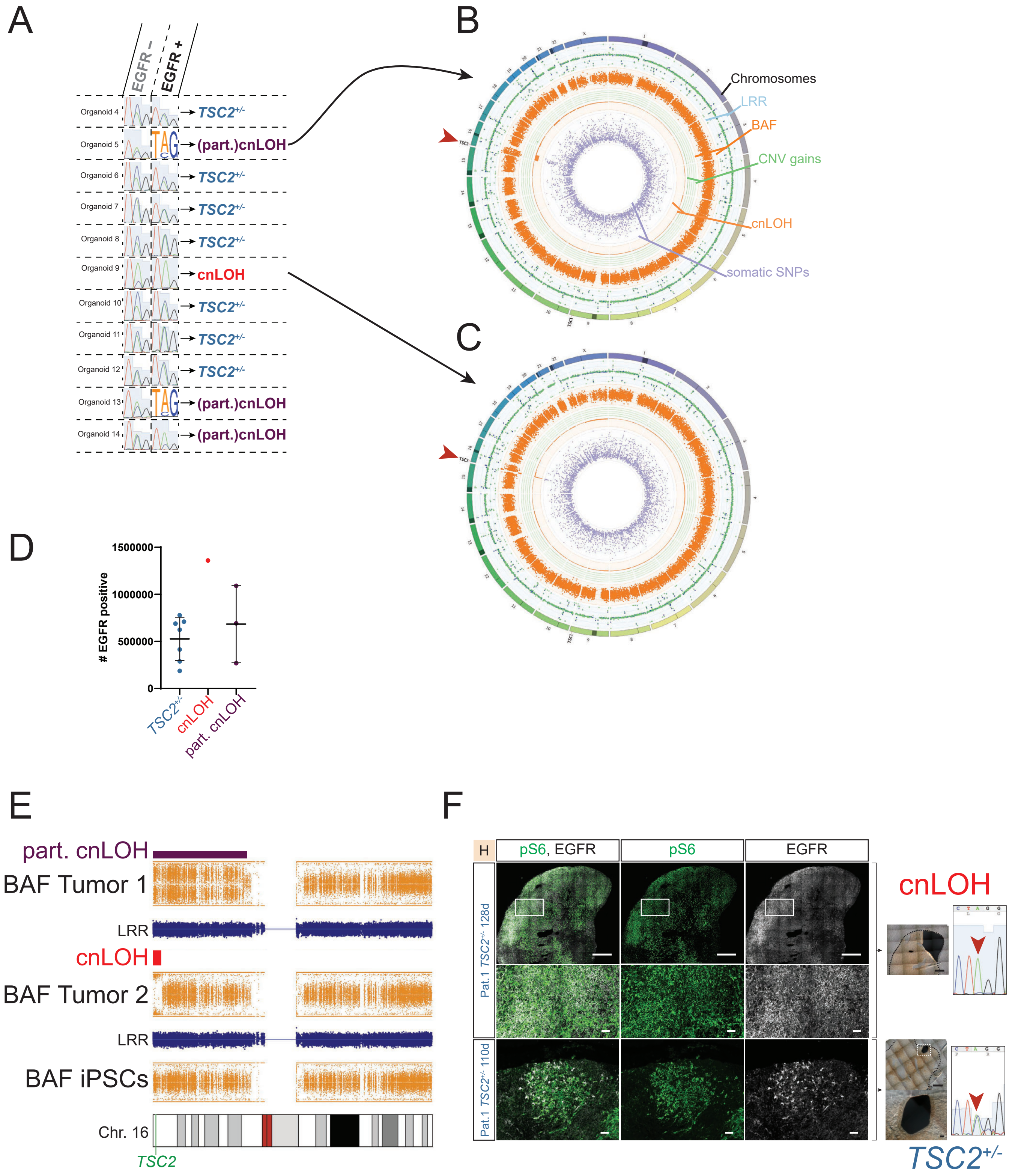
A
EGFR –
EGFR +
Organoid 4
TSC2+/-
Organoid 5
(part.)cnLOH
Organoid 6
TSC2+/-
Organoid 7
TSC2+/-
Organoid 8
TSC2+/-
Organoid 9
cnLOH
Organoid 10
TSC2+/-
Organoid 11
TSC2+/-
Organoid 12
TSC2+/-
Organoid 13
(part.)cnLOH
Organoid 14
(part.)cnLOH
B
Chromosomes
LRR
BAF
CNV gains
cnLOH
somatic SNPs
C
D
### EGFR positive
1500000
1000000
500000
0
TSC2+/-
cnLOH
part. cnLOH
E
part. cnLOH
BAF Tumor 1
LRR
cnLOH
BAF Tumor 2
LRR
BAF iPSCs
Chr. 16
TSC2
F
H
pS6, EGFR
pS6
EGFR
Pat.1 TSC2+/- 128d
Pat.1 TSC2+/- 110d
cnLOH
TSC2+/-

**Ext. Data Fig. 13 - Sequencing on TSC tumors**

**A.** Sequencing *TSC2* mutation locus of all 11 *TSC2*$^{+/-}$ -derived organoids that were genotyped after FACS. 7/11 tumors are heterozygous.

**B.** and **C.** Modified Circos Plot tumors 5 and 9. Chromosomes are depicted on outside starting with Chr. 1 clockwise. The next ring shows log-R ratio (LRR) for tumor (blue) and reference (green, *TSC2*$^{+/-}$ iPSCs are used as a reference). No aneuploidy occurs in the two tumors indicated by overlapping tumor and reference LRR in blue and green respectively. The next ring shows B-allele frequency (BAF) in orange. Only deviation of BAF from center is detected around *TSC2* locus on Chr. 16 (red arrows). Green inner ring shows inferred copy-number variation (CNV) gains. No CNV changes are detected. Inner orange circle shows predicted copy-neutral loss-of-heterozygozity (cnLOH). Percentage of cells showing cnLOH is indicated by bar towards center of the plot. Note that tumor 5 shows a bigger region of cnLOH but the percentage of cells (predicted around 50% of cells) is less than in the small region of cnLOH in tumor 9 (predicted 100% of cells). Innermost in purple somatic SNPs are shown. Only clear deviation is found around cnLOH on Chr. 16. Note that none of these parameters are changes on *TSC1* locus marked on Chr. 9.

**D.** Number of EGFR+ cells in heterozygous, complete cnLOH and partial cnLOH population shows trend towards higher numbers, indicating bigger tumors in cnLOH tumors.

**E.** More detailed view of cnLOH as in Fig. 4E. BAF shows different size of cnLOH in both tumors. Note difference in shift of BAF from midline indicating lower (Tumor 1) and higher (Tumor 2) percentage of cnLOH cells in EGFR-positive sample. As a reference *TSC2*$^{+/-}$ iPSCs are shown.

**F.** Immunostaining of consecutive slides (left) used for Laser Capture Microdissection (LCM) and corresponding sequencing shown on the right. Upper large tumor was confirmed to have cnLOH while the lower smaller tumor is still heterozygous. Note that the scale of the heterozygous tumor corresponds to the scale of the inset of the cnLOH tumor showing extreme differences in size between heterozygous and cnLOH tumors. LMD membranes with cut-out tissue section as well as Sanger Sequencing results are shown on the right.

(Scale Bars: Overview top panel F, overview LMD membrane: 500µm; Inset and bottom panel F, Inset LMD membrane: 50µm)

A

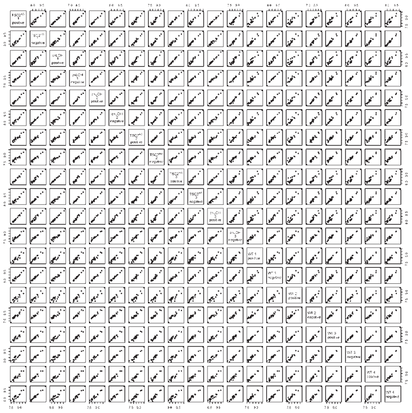

B

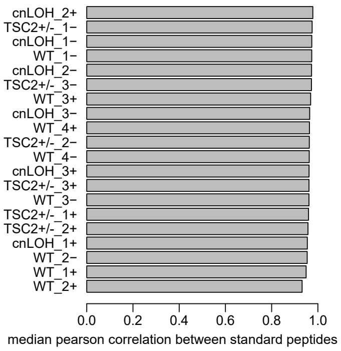


C

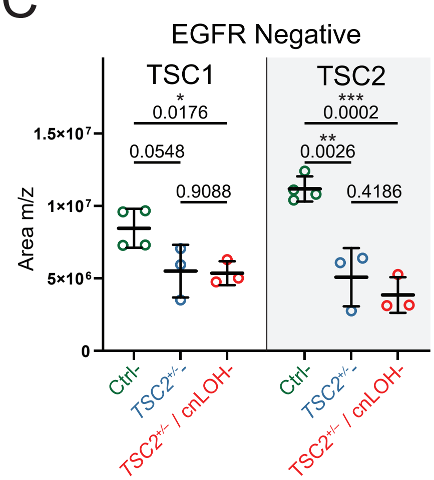


D

| protein name | gene name | database accession number | peptide sequence | m/z | charge | chemical modification | Scheduling [min] | |
|---|---|---|---|---|---|---|---|---|
| | | | | | | | Start | End |
| Hamartin | TSC1 | Q92574 | LSQAPLLPSLLK (light) | 640,4028 | 2 | | 93,56 | 97,56 |
| | | | LSQAPLLPSLLK (heavy) | 644,4099 | 2 | | 93,56 | 97,56 |
| | | | IHPELVTGSK (light) | 540,806 | 2 | | 30,71 | 34,71 |
| | | | IHPELVTGSK (heavy) | 544,8131 | 2 | | 30,71 | 34,71 |
| | | | ISLDPTEASYEDGYSVSHQISAR (light) | 842,3979 | 3 | | 77,64 | 81,64 |
| | | | ISLDPTEASYEDGYSVSHQISAR (heavy) | 845,734 | 3 | | 77,64 | 81,64 |
| | | | ELSEITTAEAEPVVPR (light) | 870,9544 | 2 | | 74,19 | 78,19 |
| | | | ELSEITTAEAEPVVPR (heavy) | 875,9585 | 2 | | 74,19 | 78,19 |
| | | | LIQQGADAHSK (light) | 584,3095 | 2 | | 13,12 | 17,12 |
| | | | LIQQGADAHSK (heavy) | 588,3166 | 2 | | 13,12 | 17,12 |
| Tuberin | TSC2 | P49815 | LLQQLQTLDSPELR (light) | 827,4621 | 2 | | 85,39 | 89,39 |
| | | | LLQQLQTLDSPELR (heavy) | 832,4663 | 2 | | 85,39 | 89,39 |
| | | | YFELVER (light) | 478,2478 | 2 | | 68,13 | 72,13 |
| | | | YFELVER (heavy) | 483,252 | 2 | | 68,13 | 72,13 |
| | | | DFVPFITK (light) | 483,7684 | 2 | | 90,23 | 94,23 |
| | | | DFVPFITK (heavy) | 487,7755 | 2 | | 90,23 | 94,23 |
| | | | SLHAEELVGR (light) | 555,7987 | 2 | | 41,08 | 45,08 |
| | | | SLHAEELVGR (heavy) | 560,8029 | 2 | | 41,08 | 45,08 |
| | | | YTEFLTGLGR (light) | 578,8035 | 2 | | 81,08 | 85,08 |
| | | | YTEFLTGLGR (heavy) | 583,8076 | 2 | | 81,08 | 85,08 |
| Signal recognition particle 14 kDa protein | SRP14 | P37108 | TSGSVYITLK (light) | 534,8004 | 2 | | 58,18 | 62,18 |
| | | | GTVEGFEPADNK (light) | 632,2962 | 2 | | 44,50 | 48,50 |
| Lamin-B1 | LMNB1 | P20700 | SLETENSALQLQVTER (light) | 909,4656 | 2 | | 72,58 | 76,58 |
| | | | LQIELGK (light) | 400,7475 | 2 | | 55,38 | 59,38 |
| | | | AEHDQLLLNYAK (light) | 707,8699 | 2 | | 58,15 | 62,15 |
| | | | QLADETLLK (light) | 515,7926 | 2 | | 59,37 | 63,37 |
| | | | C[+57,021464]QSLTEDLEFR (light) | 699,3219 | 2 | carbamidomethylation of C | 77,97 | 81,97 |
| | | | EELEQTYHAK (light) | 624,2988 | 2 | | 26,12 | 30,12 |
| | | | IESLSSQLSNLQK (light) | 723,8936 | 2 | | 72,61 | 76,61 |
| | | | IQELEDLLAK (light) | 586,3321 | 2 | | 85,49 | 89,49 |
| | | | IGDTSVSYK (light) | 485,248 | 2 | | 34,99 | 38,99 |
| Histone H1,5 | HIST1H1B | P16401 | ATGPPVSELITK (light) | 606,8454 | 2 | | 63,62 | 67,62 |
| | | | NGLSLAALK (light) | 443,7715 | 2 | | 65,90 | 69,90 |
| | | | ALAAGGYDVEK (light) | 547,2799 | 2 | | 44,62 | 48,62 |
| Glyceraldehyde-3-phosphate dehydrogenase | GAPDH | P04406 | VGVNGFGR (light) | 403,2194 | 2 | | 42,18 | 46,18 |
| | | | VGVN[+0,984016]GFGR (light) | 403,7114 | 2 | deamidation of N | 45,99 | 49,99 |
| | | | IISNASC[+57,021464]TTNC[+57,021464]LAPLAK (light) | 917,4635 | 2 | | 63,67 | 67,67 |
| | | | IISN[+0,984016]ASC[+57,021464]TTNC[+57,021464]LAPLAK (light) | 917,9555 | 2 | 2x carbamidomethylation of C | 63,65 | 67,65 |
| | | | VIPELNGK (light) | 435,2582 | 2 | carbamidomethylation of C | 44,44 | 48,44 |
| | | | VPTANVSVVDLTC[+57,021464]R (light) | 765,9009 | 2 | | 74,12 | 78,12 |
| Malate dehydrogenase, mitochondrial | MDH2 | P40926 | LTLYDIAHTPGVAADLSHIETK (light) | 789,0848 | 3 | | 95,23 | 99,23 |
| | | | GYLGPEQLPDC[+57,021464]LK (light) | 745,3714 | 2 | carbamidomethylation of C | 81,30 | 85,30 |
| | | | GC[+57,021464]DVVVIPAGVPR (light) | 669,8636 | 2 | carbamidomethylation of C | 74,25 | 78,25 |
| | | | IFGVTTLDIVR (light) | 617,3637 | 2 | | 98,31 | 102,31 |
| | | | SQETEC[+57,021464]TYFSTPLLLGK (light) | 987,4799 | 2 | carbamidomethylation of C | 99,06 | 103,06 |

**Ext. Data Fig. 14 - PRM on TSC1 and TSC2 protein levels in TSC tumors**

**A.** Pearson Correlation of all normalisation proteins (SRP14, LMNB1, H15, G3P, MDHM) across all samples showing high correlation and thus validate use for normalisation (Ctrl: N=2, n=4; *TSC2*$^{+/-}$: N=2, n=3; cnLOH: N=2, n=3; for each sample matched EGFR+ and EGFR-).

**B.** Bar graph of median pearson correlation between standard peptides as in A showing high correlation (>0.95) between normalisation peptides of samples.

**C.** Normalized levels of TSC1 and TSC2 comparing different genotypes in EGFR negative samples showing lower TSC1 and TSC2 levels in *TSC2*$^{+/-}$- and cnLOH tumor-matched EGFR- samples (unpaired t-test).

**D.** Information on all peptide fragments, as well as scheduling used in this experiment.

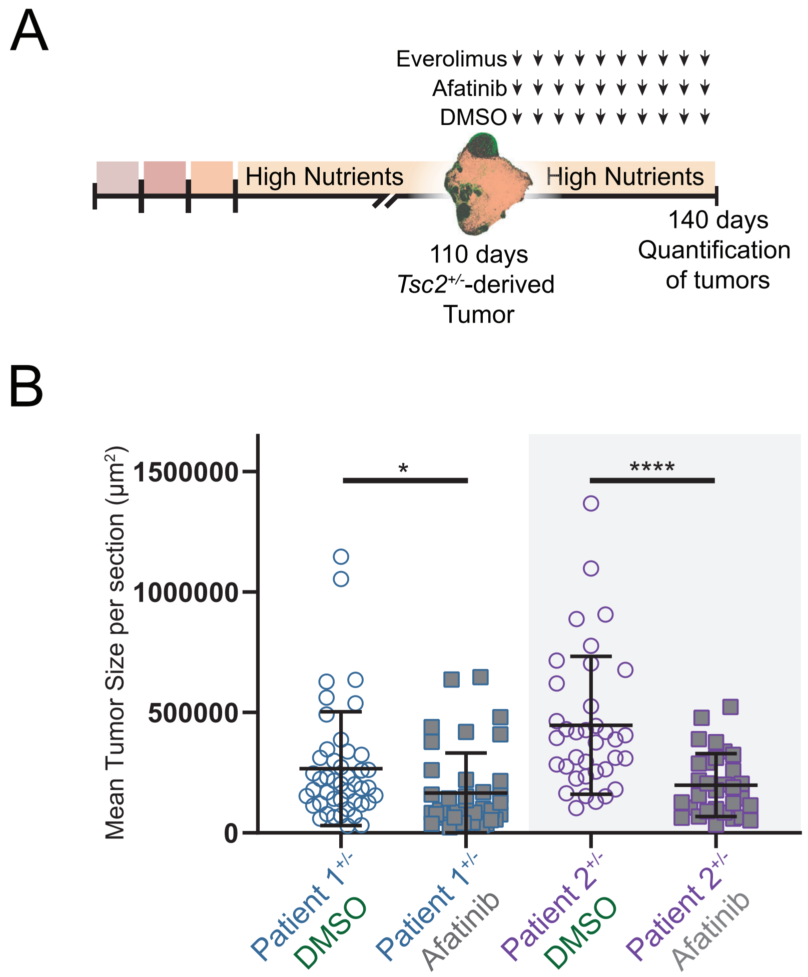


**Ext. Data Fig. 15 - Drug-testing assay**

**A.** Experimental design for tumor treatments. Organoids were grown in high-nutrient medium until 110 days after EB formation. Presence of tumors was confirmed, and organoids were split in three treatment groups. Every third day organoids were fed with H-medium supplemented with DMSO (volume adjusted to DMSO volume of other drugs), Everolimus (20nM final concentration, as previously used in 3D culture[5] or Afatinib (1µM as previously used by our lab in a glioblastoma model[6]) over the course of 30 days. Organoids were fixed and stained 140 days after EB formation.

**B.** Quantification of Mean tumor size per section shows that Afatinib significantly reduces the size of tumors (N=3, n=12; Pat.1 *TSC2*$^{+/-}$-DMSO mean=2.67E5, SD=2.34E5; Pat.1 *TSC2*$^{+/-}$-Afa mean=1.66E5, SD=1.63E5; Pat.2 *TSC2*$^{+/-}$-DMSO mean= 4.47E5, SD=2.82E5; Pat.2 *TSC2*$^{+/-}$-Afa mean=1.99E5, SD=1.28E5; Pat.1 *TSC2*$^{+/-}$-DMSO vs. Afa p=0.0266; Pat.2 *TSC2*$^{+/-}$-DMSO vs. Afa p<0.0001; Unpaired t-test).
